# Genome-wide Identification of the Genetic Basis of Amyotrophic Lateral Sclerosis

**DOI:** 10.1101/2020.11.14.382606

**Authors:** Sai Zhang, Johnathan Cooper-Knock, Annika K. Weimer, Minyi Shi, Tobias Moll, Calum Harvey, Helia Ghahremani Nezhad, John Franklin, Cleide dos Santos Souza, Cheng Wang, Jingjing Li, Eran Elhaik, Chen Eitan, Eran Hornstein, Kevin P. Kenna, Project MinE Sequencing Consortium, Jan Veldink, Laura Ferraiuolo, Pamela J. Shaw, Michael P. Snyder

## Abstract

Amyotrophic lateral sclerosis (ALS) is an archetypal complex disease centered on progressive death of motor neurons. Despite heritability estimates of 52%, GWAS studies have discovered only seven genome-wide significant hits, which are relevant to <10% of ALS patients. To increase the power of gene discovery, we integrated motor neuron functional genomics with ALS genetics in a hierarchical Bayesian model called RefMap. Comprehensive transcriptomic and epigenetic profiling of iPSC-derived motor neurons enabled RefMap to systematically fine-map genes and pathways associated with ALS. As a significant extension of the known genetic architecture of ALS, we identified a group of 690 candidate ALS genes, which is enriched with previously discovered risk genes. Extensive conservation, transcriptome and network analyses demonstrated the functional significance of these candidate genes in motor neurons and disease progression. In particular, we observed a genetic convergence on the distal axon, which supports the prevailing view of ALS as a distal axonopathy. Of the new ALS genes we discovered, we further characterized *KANK1* that is enriched with coding and noncoding, common and rare ALS-associated genetic variation. Modelling patient mutations in human neurons reduced *KANK1* expression and produced neurotoxicity with disruption of the distal axon. RefMap can be applied broadly to increase the discovery power in genetic association studies of human complex traits and diseases.

## INTRODUCTION

ALS is an untreatable, universally fatal and relatively common neurodegenerative disease with a lifetime risk of ~1/350 in the UK. The hallmark of the disease is motor neuron loss leading to respiratory failure and death (Hardiman et al., 2017). 10% of ALS is autosomal dominant, and even for sporadic ALS (sALS), the heritability is estimated to be ~50% (Ryan et al., 2019; Trabjerg et al., 2020). Genome-wide association studies (GWAS) in ALS (Nicolas et al., 2018; van Rheenen et al., 2016) have identified seven genome-wide significant loci, which have been linked to missense mutations. However, these changes occur in <10% of ALS patients, so there are likely to be a large number of missing ALS risk genes.

ALS GWAS studies to date have lost power by considering genetic variants in isolation, whereas in reality, a biological system is the product of a large number of interacting partners (Li et al., 2019; Wang et al., 2011). Moreover, noncoding regulatory regions of the genome have been relatively neglected in efforts to pinpoint the genetic basis of ALS, despite their functional synergy with the coding sequence (Cooper-Knock et al., 2020; Wang et al., 2018). Indeed, GWAS studies have suggested that a significant proportion of missing heritability in ALS is distributed throughout noncoding chromosomal regions (Nicolas et al., 2018; van Rheenen et al., 2016). The function of noncoding DNA is often tissue, disease, or even cell-type specific (Heinz et al., 2015), and the understanding of the cell-type-specific biological function in complex neurological diseases has been improving (Bryois et al., 2020; Lopategui Cabezas et al., 2014). This therefore creates an opportunity to dramatically reduce the search space and so boost the power to discover ALS genetic risk, by focusing on genomic regions that are functional within the cell type of interest, i.e., motor neurons (MNs) (Cooper-Knock et al., 2013).

Here, we present RefMap (**<u>Re</u>**gional **<u>F</u>**ine-**<u>map</u>**ping), a hierarchical Bayesian model to perform genome-wide identification of disease-associated genetic variation within active genomic regions. RefMap utilizes cell-type-specific epigenetic profiling to determine the prior probability of disease-association for each region. This reduces the search space by >90% given that a limited proportion of the genome is active in any specific cell type. ALS is notable for the selective vulnerability of MNs (Cooper-Knock et al., 2013). However, MNs are difficult to study in post-mortem tissues (Corces et al., 2020) because of their relative sparsity, so a different approach is needed. We performed exhaustive transcriptomic and epigenetic profiling, including RNA-seq, ATAC-seq, histone ChIP-seq and Hi-C, for motor neurons derived from fibroblasts of neurologically normal controls. We hypothesized that the genetic variation within regulatory regions may alter the expression of their target genes, and we proposed that disease-associated variants are likely to reduce gene expression via interfering with regulation. Applying RefMap to perform genome-wide fine-mapping based on ALS GWAS data (**Fig. 1a**) identified 690 ALS-associated genes, including previous GWAS hits and even known ALS genes not previously detected in GWAS studies.

**Figure 1.**
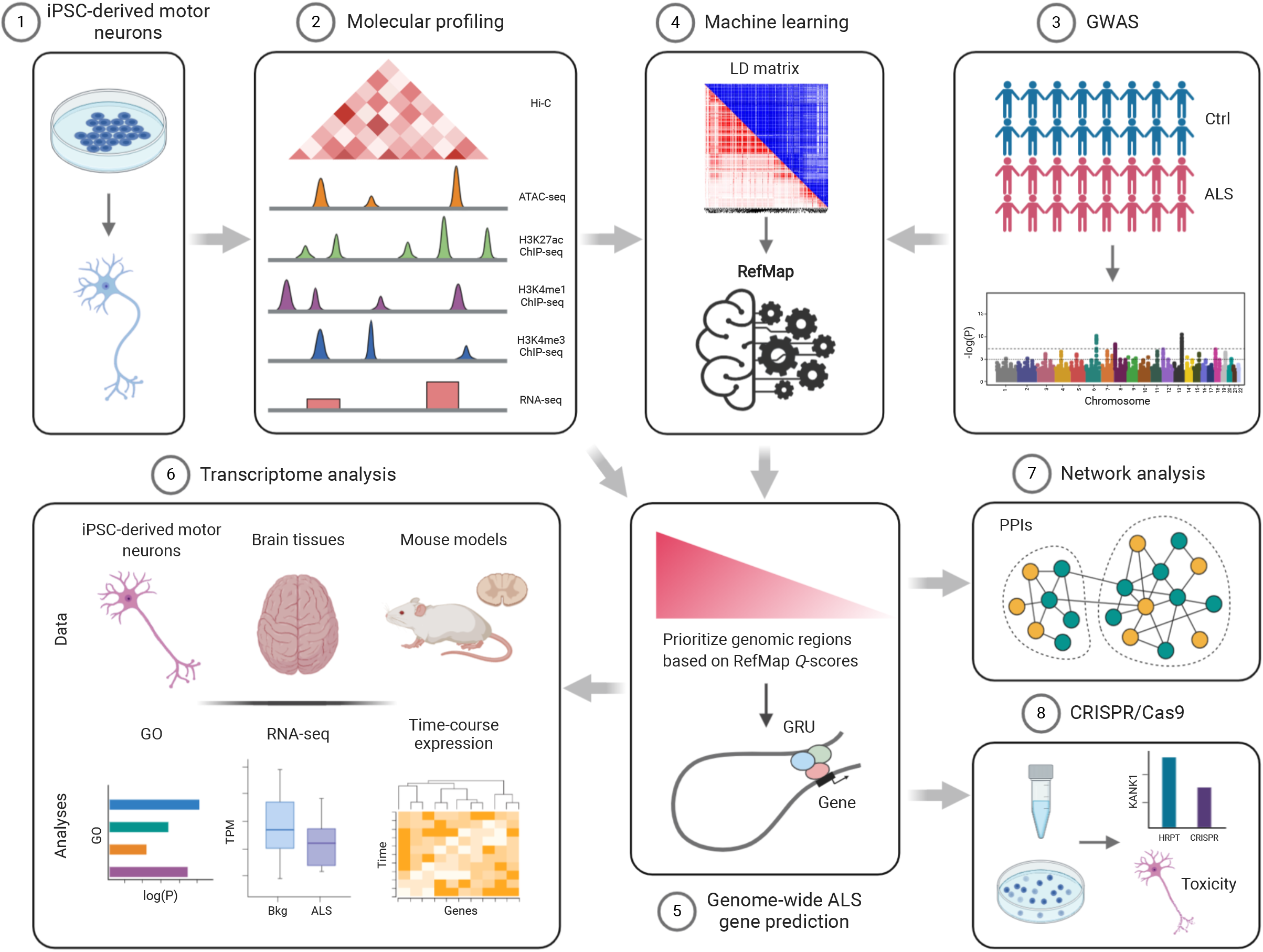
Genome-wide identification of ALS-associated genes. (**a**) Schematic of the study design for identifying ALS risk genes by integrating ALS genetics with functional genomics from motor neurons. (1-2) We sequenced the transcriptome and epigenome of the iPSC-derived motor neurons. By integrating (3) ALS genetics with functional genomics of MNs, (4) a machine learning model called RefMap was developed to fine-map ALS-associated regions. (5) After mapping those identified regions to their target genes, 690 ALS-associated genes were pinpointed. (6) Transcriptome analysis based on iPSC-derived MNs, human tissues and mouse models, as well as (7) network analysis were performed to demonstrate the functional significance of RefMap genes in ALS. (8) CRISPR/Cas9 reproduction of newly identified ALS-associated *KANK1* mutations experimentally verified the proposed link to neuronal toxicity. The LD heatmap matrix in (4) was visualized in both *R^2^* (red) and *D’* (blue) using LDmatrix (https://ldlink.nci.nih.gov/?tab=ldmatrix). GRU=Gene Regulatory Unit; GO=Gene Ontology. (**b**) Epigenetic profiling data from motor neurons is internally consistent. Markers of genomic activity are significantly enriched in promoter regions of high-expressed genes compared to low-expressed genes. Circle area is proportional to % overlap. Hi-C data was scaled by a factor of ten for clarity. (**c**) Graphical representation of RefMap. Observed variables were annotated in grey, local hidden variables were in green and global latent variables were in pink (**Methods**). (**d**) A region (chr12:112,036,001-112,038,000) around *ATXN2* was precisely pinpointed by RefMap because of elevated SNP *Z*-scores and enriched epigenome peaks (ATAC-seq, H3K27ac and H3K4me3 histone ChIP-seq). The output of RefMap was labeled as *Q*-score. ATAC-seq and ChIP-seq signals were shown in fold change (FC) based on one replicate from sample CS14.

We explored the functional significance of RefMap ALS genes based on a series of orthogonal analyses. Population genetics revealed that RefMap genes consist of conserved sequences, suggesting that their functions are important and not subject to genetic redundancy. Transcriptome data from MNs, human tissues and mouse models demonstrated that RefMap genes are down-regulated in ALS patients, consistent with our aforementioned hypothesis. Network analysis of protein-protein interactions (PPIs) identified two modules enriched with RefMap genes. These modules are enriched with biological functions localized to the distal axon of MNs, suggesting that neurotoxicity may be initiated in this subcompartment, which is consistent with previous literature (Frey et al., 2000; Moloney et al., 2014). Finally, we have further characterized a new ALS gene, i.e., *KANK1.* Common and rare genetic variants that alter *KANK1* expression were shown to be associated with ALS and neuronal toxicity. RefMap provides a promising framework to pinpoint the genetic bases of human complex traits and diseases based on GWAS data.

## RESULTS

### Transcriptomic and epigenetic profiling of iPSC-derived motor neurons

To identify genomic regions key to motor neuron function, we performed transcriptomic and epigenetic profiling of iPSC-derived motor neurons from neurologically normal individuals (**Supplementary Fig. 1**). The cells exhibited homogenous expression of the lower motor neuron markers, including TUJ1, Chat, SMI, MAP2 and NeuN (**Supplementary Fig. 1a**). We prepared RNA-seq (Wang et al., 2009), ATAC-seq (Buenrostro et al., 2015), H3K27ac, H3K4me1 and H3K4me3 ChIP-seq (Creyghton et al., 2010), as well as Hi-C (van Berkum et al., 2010) libraries using two technical replicates and three biological replicates per assay. Sequencing data were processed and quality control (QC) was performed according to the ENCODE 4 standards (ENCODE Project Consortium et al., 2020), and all samples exceeded ENCODE standard QC measures (**Supplementary Tables 1-4**).

ATAC-seq identifies open and functional chromatin regions, which is complementary to the profiling of transcript expression by RNA-seq. H3K27ac, H3K4me1 and H3K4me3 ChIP-seq assays pinpoint active enhancer (Pennacchio et al., 2013) regions, which are important noncoding regions for the regulation of gene expression. Hi-C profiling of threedimensional (3D) genome structure is essential to map regulatory regions including enhancers, to their target genes. Our MN epigenetic profiling successfully reduced the search space for ALS-associated genetic variation by >90%. Specifically, total ATAC-seq peak regions across all biological replicates covered 4.9% of the genome.

To measure the consistency between distinct motor neuron profiles, we used our RNA-seq dataset to identify promoter regions for high (>90th centile) and low (<10th centile) expressed transcripts. We compared enrichment of ATAC-seq and histone ChIP-seq peak regions, and Hi-C loops in high versus low expressed promoters. Significant enrichment within highly expressed promoters was confirmed for ATAC-seq (*P*=1.1e-182, odds ratio (OR)=1.9, Fisher’s exact test), H3K27ac ChIP-seq (*P*=2.0e-57, OR=2.2, Fisher’s exact test), H3K4me1 ChIP-seq (*P*=8.5e-57, OR=1.9, Fisher’s exact test), H3K4me3 ChIP-seq (*P*=4.8e-196, OR=2.6, Fisher’s exact test), and Hi-C loops (*P*=4.0e-14, OR=1.3, Fisher’s exact test) (**Fig. 1b**). Similarly, epigenetic peak regions were enriched in MN Hi-C loops: ATAC-seq (*P*<1.0e-198, OR=1.9, Fisher’s exact test), H3K27ac ChIP-seq (*P*<1.0e-198, OR=2.0, Fisher’s exact test), H3K4me1 ChIP-seq (*P*<1.0e-198, OR=2.0, Fisher’s exact test), and H3K4me3 ChIP-seq (*P*<1.0e-198, OR=1.7, Fisher’s exact test). These observations confirm that our epigenetic profiling captured functionally significant genomic variation, and that our epigenetic profiles were internally consistent.

### RefMap identifies ALS risk genes

Mismatch between the relatively small number of characterized ALS risk genes and the estimate of high heritability suggests that a new approach is required to discover more ALS-associated genetic variation. Here, we designed a hierarchical Bayesian network named RefMap that exploits the epigenetic profiling of MNs to reduce the search space and so improve the statistical power to discover ALS-associated loci across the genome. Specifically, RefMap integrates the prior probability of significance derived from the epigenome of MNs, with allele effect sizes estimated from GWAS (**Figs. 1a** and **1c**, **Methods**). Based on a linear genotype-phenotype model (**Supplementary Notes**), RefMap first disentangles effect sizes from GWAS *Z*-scores, which are confounded by the structure of linkage disequilibrium (LD). Effect sizes are then summarized across genomic regions in individual LD blocks. Those regions that are within active chromatin, and where the distributions of allele effect sizes are shifted from the null distribution, are prioritized by the algorithm (**Methods**).

In our study, the *Z*-scores were calculated based on the largest published ALS GWAS study (Nicolas et al., 2018; van Rheenen et al., 2016), including genotyping of 12,577 sporadic ALS patients and 23,475 controls. An epigenetic signal was calculated from a linear combination of MN chromatin accessibility and histone marks specific to active enhancer regions (**Methods**). We defined LD blocks as 1Mb windows, where we assumed significant internal LD but negligible external LD (Loh et al., 2015). Within LD blocks, SNP correlations were estimated based on the European population (EUR) data from the 1000 Genomes Project (Consortium and The 1000 Genomes Project Consortium, 2015). With this information, RefMap scanned the genome in 1kb windows and identified all regions that are likely to harbor ALS-associated genetic variation (**Figs. 1c** and **1d**, **Methods**, and **Supplementary Table 5**).

Next, we mapped ALS-associated regions identified by RefMap to expressed transcripts in MNs (RNA-seq, TPM>=1), based on their regulation targets. We defined regulation targets as genes that overlap either ALS-associated regions by extension, or via their Hi-C loop anchors (**Methods**). This resulted in 690 ALS-associated genes (**Supplementary Table 6**). Among this list, we discovered well-known ALS genes, including *C9orf72 (DeJesus-Hernandez et al., 2011)* and *ATXN2 (Elden et al., 2010)* (**Fig. 1d**). Indeed, RefMap genes are enriched with an independently curated list (**Supplementary Table 7**) of ALS genes including previous GWAS hits (*P*=5.20e-3, OR=2.07, Fisher’s exact test) and also with clinically reportable (ClinVar(Landrum et al., 2018)) ALS genes (*P*=0.03, OR=3.06, Fisher’s exact test). Interestingly, certain ALS genes, such as *UNC13A (Daoud et al., 2010; Diekstra et al., 2012),* are missing from RefMap genes, but their paralogues are present, including *UNC13B,* which is consistent with a functional overlap. If we consider paralogues as equivalent to ALS genes, then the enrichment of RefMap genes with known ALS genes is further increased (curated: *P*=6.12e-43, OR=8.71; ClinVar: *P*=6.40e-14, OR=12.26; Fisher’s exact test).

As a negative control, we randomly shuffled SNP *Z*-scores, in which case there was no overlap between RefMap outputs and known ALS genes. Additional shuffling of epigenetic features disrupted the signal further such that there were no significant RefMap outputs. This illustrates the dependence of RefMap on the two primary inputs: GWAS *Z*-scores and MN epigenetic features.

As a comparison to RefMap, we also applied MAGMA (v1.08) (de Leeuw et al., 2015), Pascal (Lamparter et al., 2016) and PAINTOR (Kichaev et al., 2014), which are three of the most popular methods for integrative analysis based on GWAS summary statistics. After multiple testing correction, MAGMA identified 10 genes as ALS-associated (*P*<2.76×10^−6^), and Pascal identified 5 genes (P<2.29×10^−6^), both including the known ALS gene *C9orf72.* Unlike MAGMA and Pascal, PAINTOR includes the capacity to integrate epigenetic annotations. Despite this, PAINTOR pinpointed only two ALS-associated genes: *MOB3B* and *LOC105376001* (**Supplementary Table 8**). This exercise indicates that MAGMA, Pascal and PAINTOR do not substantially address the problem of missing heritability in ALS and demonstrates very effectively the significant statistical advantage offered by our Bayesian approach.

### Conservation analysis demonstrates the functional importance of RefMap genes

A large proportion of RefMap ALS genes were identified because of ALS-associated genetic variation within noncoding regulatory regions. We hypothesized that the functional consequence of pathogenic genetic variation within regulatory regions is likely to be reduced expression of the target genes. A conservation analysis was first carried out, revealing that change in the expression of RefMap ALS genes is likely to be pathogenic based on population genetics.

Conservation refers to DNA sequences that are preserved in the population presumably because disruption would be deleterious. Conservation can be quantified by the haploinsufficiency (HI) score, which is a measure of functional similarity to known haplosufficient and haploinsufficient genes (Huang et al., 2010). Conservation is also related to intolerance scores, in which the rate of observed mutation of a gene in the population is compared to the expected rate in the absence of negative selection (Fadista et al., 2017; Karczewski et al., 2020; Petrovski et al., 2013). In particular, a lower than expected mutation rate implies intolerance to mutation. We discovered that RefMap genes are significantly haploinsufficient based on their HI score (*P*=2.59e-19, one-sided Wilcoxon rank-sum test; **Fig. 2a**), and intolerant to loss of function mutations within the Exome Aggregation Consortium (ExAC) dataset (Lek et al., 2016) as measured by LoFtool score (Fadista et al., 2017): *P*=2.28e-4 (one-sided Wilcoxon rank-sum test; **Fig. 2b**). They are also intolerant to other mutation types as measured by RVIS score (Petrovski et al., 2013): *P*=8.08e-13 (one-sided Wilcoxon rank-sum test; **Fig. 2c**), as well as within the larger gnomAD (v.2.1) dataset as measured by o/e score (Karczewski et al., 2020): *P*=4.08e-10 (one-sided Wilcoxon rank-sum test; **Fig. 2d**). Taken together, these results support the functional significance of RefMap ALS genes.

**Figure 2.**
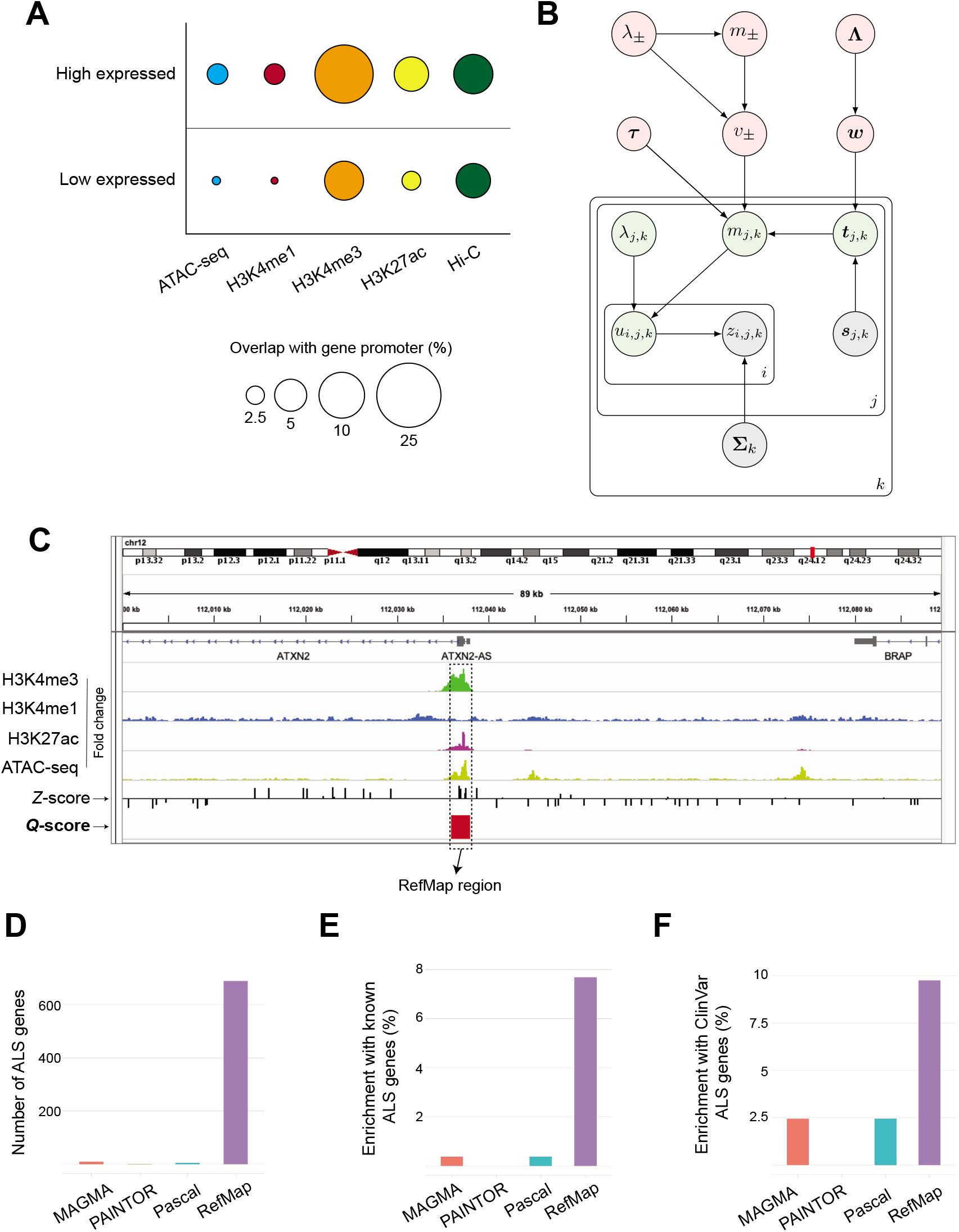
RefMap genes are intolerant to loss of function. (**a-d**) Comparison of (**a**) haploinsufficiency score, (**b**) LoFtool percentile, (**c**) RVIS-ExAC percentile and (**d**) o/e score between RefMap genes and all the genes in the transcriptome. All comparisons were performed using the one-sided Wilcoxon rank-sum test. RefMap genes showed a significant increase in HI score (**a**) and a decrease in LoFtool percentile (**b**), RVIS-ExAC percentile (**c**) and o/e score (**d**). The bottom and top of the boxes indicate the first and third quartiles, respectively, where the black line in between indicates the median. The whiskers denote the minimal value within 1.5 interquartile range (IQR) of the lower quartile and the maximum value within 1.5 IQR of the upper quartile. The plus symbols represent outliers. In **e**, the black dashed lines indicate the lower and upper limits of the regions with regular scale. Outliers outside of the black dashed lines are visualized with compressed scale in regions surrounded by gray lines for better visualization.

### Transcriptome analysis supports functional significance of RefMap genes in motor neurons and in ALS

We have hypothesized that the ALS-associated genetic variation identified by RefMap is likely to be pathogenic through altered expression of the 690 RefMap genes. We have also demonstrated, based on population genetics, that the function of RefMap genes is highly sensitive to changes in expression. To explore this possibility further, we examined whether change in the expression of RefMap genes is associated with ALS, using transcriptome data from patient-derived MNs, central nervous system (CNS) tissues and an ALS animal model.

First, we inspected the expression of RefMap genes in our iPSC-derived MNs from neurologically normal individuals. RefMap genes were upregulated (*P*=3.07e-17, onesided Wilcoxon rank-sum test; **Fig. 3a**) compared to the overall transcriptome, indicating their importance in normal MN function. No differential expression was observed for genes derived from RefMap using randomly shuffled *Z*-scores.

**Figure 3.**
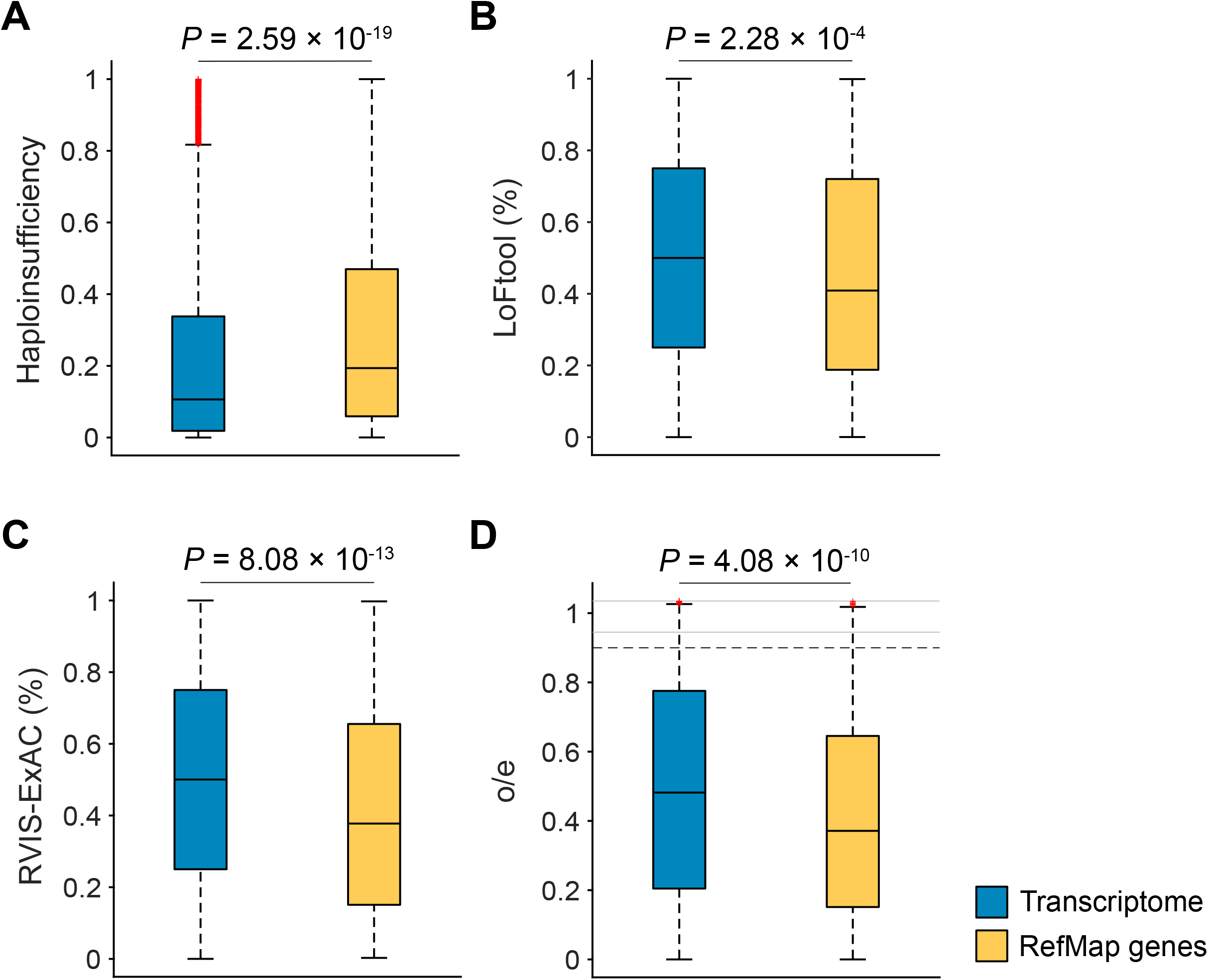
Transcriptomics supports the functional importance of RefMap genes in motor neurons and in ALS. (**a**) RefMap genes were upregulated compared to the transcriptome in iPSC-derived motor neurons from neurologically normal individuals (n=3). For a fair comparison, we only considered those genes with expressed transcripts (TPM>=1) in the transcriptome, following a similar procedure in mapping ALS-associated regions to their targets. (**b**) RefMap genes were downregulated in post-mortem CNS tissue from *C9orf72*-ALS (n=8) and sporadic ALS (n=10) patients compared to neurologically normal controls (n=17). FC=Frontal Cortex; CB=Cerebellum. (**c**) RefMap genes were downregulated in iPSC-derived motor neurons from ALS patients (n=55) compared to neurologically normal controls (n=15). All comparisons in **a-c** were performed using the one-sided Wilcoxon rank-sum test, and the Benjamini-Hochberg (BH) correction was carried out in **b**. In **a-c**, the bottom and top of the boxes indicate the first and third quartiles, respectively, where the black line in between indicates the median. The whiskers denote the minimal value within 1.5 IQR of the lower quartile and the maximum value within 1.5 IQR of the upper quartile. The plus symbols represent outliers. In **b**, the black dashed lines indicate the lower and upper limits of the regions with regular scale. Outliers outside of the black dashed lines are visualized with compressed scale in regions surrounded by gray lines for better visualization. (**d**) Hierarchical clustering of expression changes of RefMap genes during disease progression based on the *SOD1*-G93A mouse model. RefMap genes were mapped to their mouse homologs (n=510). Gene expression levels were estimated using the β scores calculated in (Maniatis et al., 2019), and were averaged across different sections of spinal cords at each time point. Time points p30, p70, p100, and p120 represent presymptomatic, onset, symptomatic and end-stage, respectively. Difference of gene expressions between *SOD1*-G93A and *SOD1*-WT mice at each time point was quantified by the difference of corresponding β (Δβ). Before clustering, Δβ were standardized across genes, and one minus correlation was used as the clustering distance. (**e**) Two distinct expression patterns (C1: 286 genes; C2: 224 genes) of RefMap genes identified after clustering. The larger cluster C1 was progressively downregulated during ALS progression. Solid plot represents the mean of expressions within each cluster, and the standard error was shown as shading. (**f**) Gene ontology analysis of C1, showing that C1 is enriched with functions related to the motor neuron distal axon and synapse. GOBP=Gene Ontology Biological Process; GOCC=Gene Ontology Cellular Compartment. Dashed line represents *P*=0.05.

Next, we examined the expression of RefMap ALS genes in CNS tissues derived from ALS patients (n=18) and controls (n=17) (Prudencio et al., 2015). We hypothesized that RefMap genes would be downregulated in ALS patient tissues. As expected, a significant decrease in the expression of RefMap genes was observed in both frontal cortex *(C9orf72-ALS* (cALS): false discovery rate (FDR)=0.002, one-sided Wilcoxon rank-sum test) and cerebellum (*C9orf72-*ALS: FDR=0.002; sporadic ALS: FDR=0.005) of ALS patients compared to the overall transcriptome (**Fig. 3b**). As an independent validation, we analyzed gene expression within iPSC-derived MNs from ALS patients (n=55, https://www.answerals.org/), and confirmed that RefMap genes were downregulated (*P*=3.85e-04, one-sided Wilcoxon rank-sum test; **Fig. 3c**) compared to neurologically normal controls (n=15).

Finally, we used longitudinal data to infer whether changes in expression of RefMap ALS genes occur upstream or downstream in the development of neuronal toxicity. To achieve this, we utilized the *SOD1-*G93A-ALS mouse model, which is the best characterized ALS model to date (Philips and Rothstein, 2015) and the only model featuring consistent and reproducible loss of spinal cord MNs that mirrors the human disease. We examined longitudinal gene expression averaged across spinal cord sections from *SOD1*-G93A (n=32) and *SOD1*-WT (n=24) mice (Maniatis et al., 2019). Four time points were sampled, including presymptomatic (p30), onset (p70), symptomatic (p100) and end-stage (p120). The model-estimated expression levels (*β*) (Maniatis et al., 2019) were adopted to quantify the gene expression difference (*Δβ*) between diseased and control mice at different time points. To determine the expression changes of RefMap genes over the course of ALS pathogenesis, we first mapped RefMap genes to their mouse homologs (n=510), and then performed unsupervised clustering on gene expressions over time. We identified two different expression patterns for RefMap homologs (**Figs. 3d** and **3e**) with verified clustering quality (**Supplementary Fig. 2**). Strikingly, the largest group (286/510) of RefMap homologs were progressively downregulated through consecutive disease stages (C1; **Figs. 3d** and **3e**, **Supplementary Table 9**), consistent with our human observations. Functional enrichment analysis (Kuleshov et al., 2016) of C1 genes revealed significant enrichment with functions associated with motor neuron biology (**Fig. 3f**), including ‘cholinergic synapse’, ‘axon’ and ‘cytoskeleton’, which is consistent with known ALS biology (Cooper-Knock et al., 2013) and with the prevailing view of ALS as a distal axonopathy (Frey et al., 2000; Moloney et al., 2014). C2 genes do not contain significant functional enrichment (data not shown).

### Systems analysis dissect ALS-associated functional modules

We have used RefMap to extend the number of ALS-associated risk genes to 690. We aimed to assess whether these genes are functionally consistent with current knowledge regarding the biology of MNs and ALS. Genes do not function in isolation and therefore, rather than examining individual genes, we mapped RefMap ALS genes to the global protein-protein interaction (PPI) network and inspected functional enrichment of ALS-associated network modules.

We first extracted high-confidence (combined score >700) PPIs from STRING v11.0(Szklarczyk et al., 2019), which include 17,161 proteins and 839,522 protein interactions. To eliminate the bias of hub genes(Krishnan et al., 2016), we performed the random walk with restart algorithm over the raw PPI network to construct a smoothed network based on those edges with weights in the top 5% (**Supplementary Table 10**, **Methods**). Next, this smoothed PPI network was decomposed into non-overlapping subnetworks using the Louvain algorithm(Blondel et al., 2008a) that maximizes the modularity to detect communities from a network. This process yielded 912 different modules (**Supplementary Table 11**), in which genes within modules were densely connected with each other but sparsely connected with genes in other modules. As a negative control, we constructed 100 shuffled networks by randomly rewiring the PPI network while keeping the same number of neighbors. None of the randomized networks achieved the same modularity of our smoothed network after clustering, demonstrating the significance of our derived gene modules (*P*<0.01; **Supplementary Fig. 3a**).

RefMap ALS genes were then mapped to individual modules, and two modules were found to be significantly enriched with RefMap genes: M421 (721 genes; FDR<0.1, hypergeometric test; **Fig. 4a**) and M604 (308 genes; FDR<0.1, hypergeometric test; **Fig. 4b**) (**Supplementary Table 11**). Functionally M421 is enriched with GO/KEGG terms related to the distal axon, including synapse and axonal function within motor neurons (**Fig. 4c**). M421 is also enriched with genes related to relevant neurodegenerative diseases, including ‘amyotrophic lateral sclerosis’ and ‘Alzheimer’s disease’. M604 is enriched with GO/KEGG terms related to the actin cytoskeleton and axonal function (**Fig. 4d**). Notably, the actin cytoskeleton is key for neuronal function and for axonal function in particular. Overall, the functional enrichment of both modules highlights an important role of the distal axon in ALS etiology (**Fig. 4e**), which is consistent with previous literature (Frey et al., 2000; Moloney et al., 2014). Finally, both M421 and M604 were overexpressed in control iPSC-derived MNs (**Fig. 4f**), in a similar manner to the total set of RefMap genes. Interestingly, many functions ascribed to M421 and M604 overlap with the functions of the C1 cluster from our analysis of the *SOD1-G93A* mouse model (**Fig. 3f**), demonstrating a functional convergence of RefMap ALS genes.

**Figure 4.**
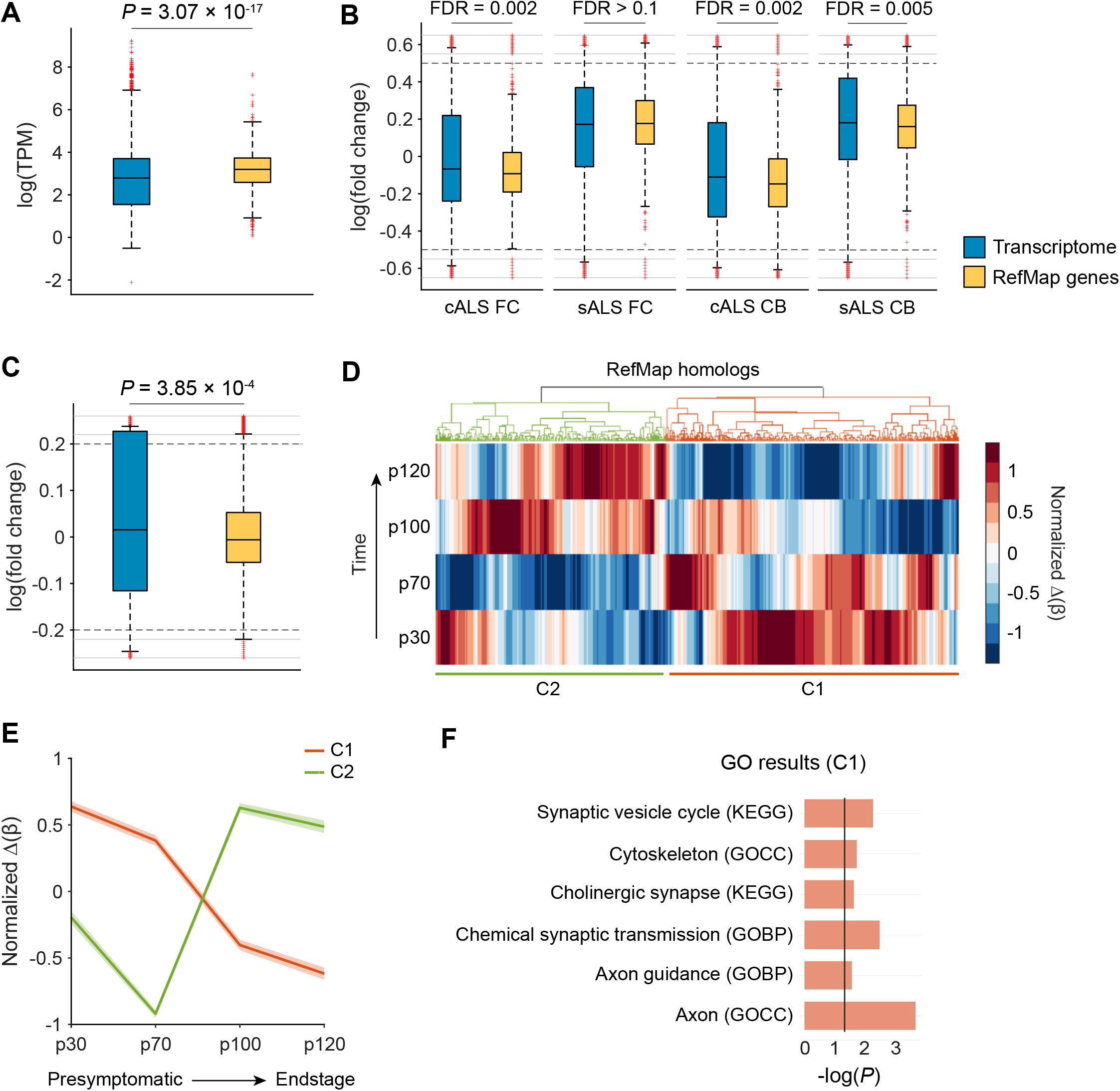
Protein-protein interaction network analyses associate RefMap genes with distal axonopathy within motor neurons. (**a-b**) PPI network analysis revealed two modules that are significantly (FDR<0.1) enriched with RefMap genes: (**a**) M421 (721 genes) and (**b**) M604 (308 genes). Hypergeometric test was performed to quantify the enrichment followed by the BH correction. Module nodes were colored to demonstrate RefMap enrichment, where RefMap genes are in blue and other module genes are in yellow. Edge thickness is proportional to STRING confidence score (>700). (**c-d**) RefMap modules, including (**c**) M421 and (**d**) M604, are enriched for motor neuron functions localized within the distal axon. GOBP=Gene Ontology Biological Process; GOCC=Gene Ontology Cellular Compartment. Dashed line represents *P*=0.05. (**e**) Representation of pathways enriched in each module (**c** and **d**) in MNs. (**f**) RefMap modules were highly expressed within control motor neurons, consistent with an important role in motor neuron function. All comparisons were performed using the one-sided Wilcoxon rank-sum test. The bottom and top of the boxes indicate the first and third quartiles, respectively, where the black line in between indicates the median. The whiskers denote the minimal value within 1.5 IQR of the lower quartile and the maximum value within 1.5 IQR of the upper quartile. The plus symbols represent outliers. The black dashed lines indicate the lower and upper limits of the regions with regular scale. Outliers outside of the black dashed lines are visualized with compressed scale in regions surrounded by gray lines for better visualization.

### Rare variant burden analysis is consistent with KANK1 as a novel ALS risk gene

Among all ALS-associated active regions identified by RefMap, chr9:663,001-664,000 has the highest concentration of ALS risk SNPs (22 SNPs). This region lies within intron 2 of *KANK1* and consists of independently annotated ENSEMBL regulatory features, including an enhancer element (ENSR00000873709) and a CTCF binding site (ENSR00000873710) (**Fig. 5a**). Overlap with independently annotated features supports the utility of RefMap to identify functional regulatory regions within noncoding DNA. We hypothesized that ALS-associated genetic variation within chr9:663,001-664,000 would reduce the expression of *KANK1,* leading to MN toxicity. Existing biological characterization of *KANK1* is consistent with our hypothesis: *KANK1* is expressed in motor neurons, functions in actin polymerization and deletion of this gene results in a severe developmental phenotype with MN loss (Lerer et al., 2005).

**Figure 5.**
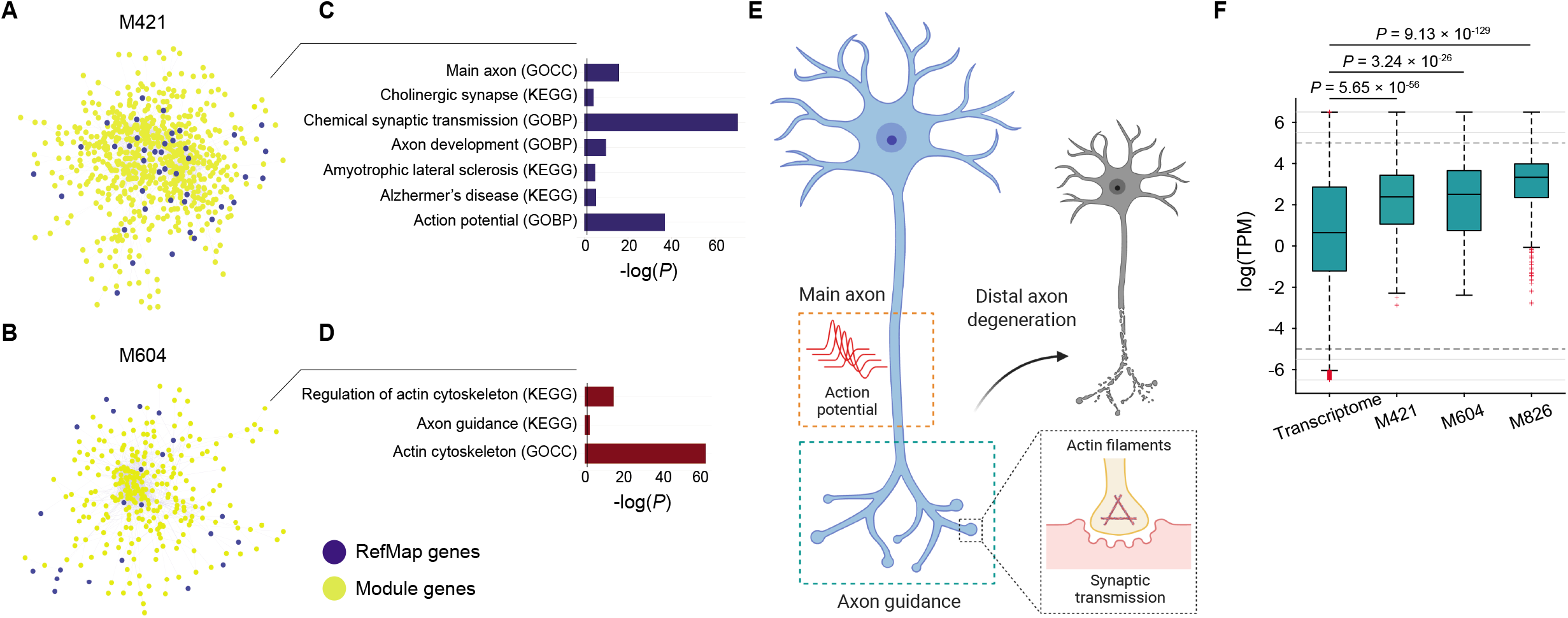
Reduced *KANK1* expression is associated with ALS-associated genetic variants and produces neurotoxicity. (**a**) We identified a high density of ALS-associated genetic variants within a region at chr9:663001-664000, which overlaps with the regulatory regions in iPSC-derived control motor neurons as well as in the ENSEMBL regulatory build. (**b**) Whole genome sequencing data from sporadic ALS patients (n=5,594) and neurologically normal controls (2,238) was analyzed to determine the frequency of rare deleterious variants within *KANK1* coding and regulatory sequences. ALS-associated rare variants are shown. All variants were present in a single patient unless stated. No variant was found in a control individual. (**c**) To experimentally evaluate ALS-associated *KANK1* variants, we performed CRISPR/Cas9 perturbation proximate to patient mutations in enhancer and coding regions within SH-SY5Y neurons, including resection of the chr9:663001-664000 region. Edited neurons revealed (**d**) reduced viability, (**e**) reduced axonal length and (**f**) reduced axonal-branch length compared to HPRT-edited controls. Data shown is mean and standard deviation. Neuronal viability was quantified relative to HPRT-edited controls.

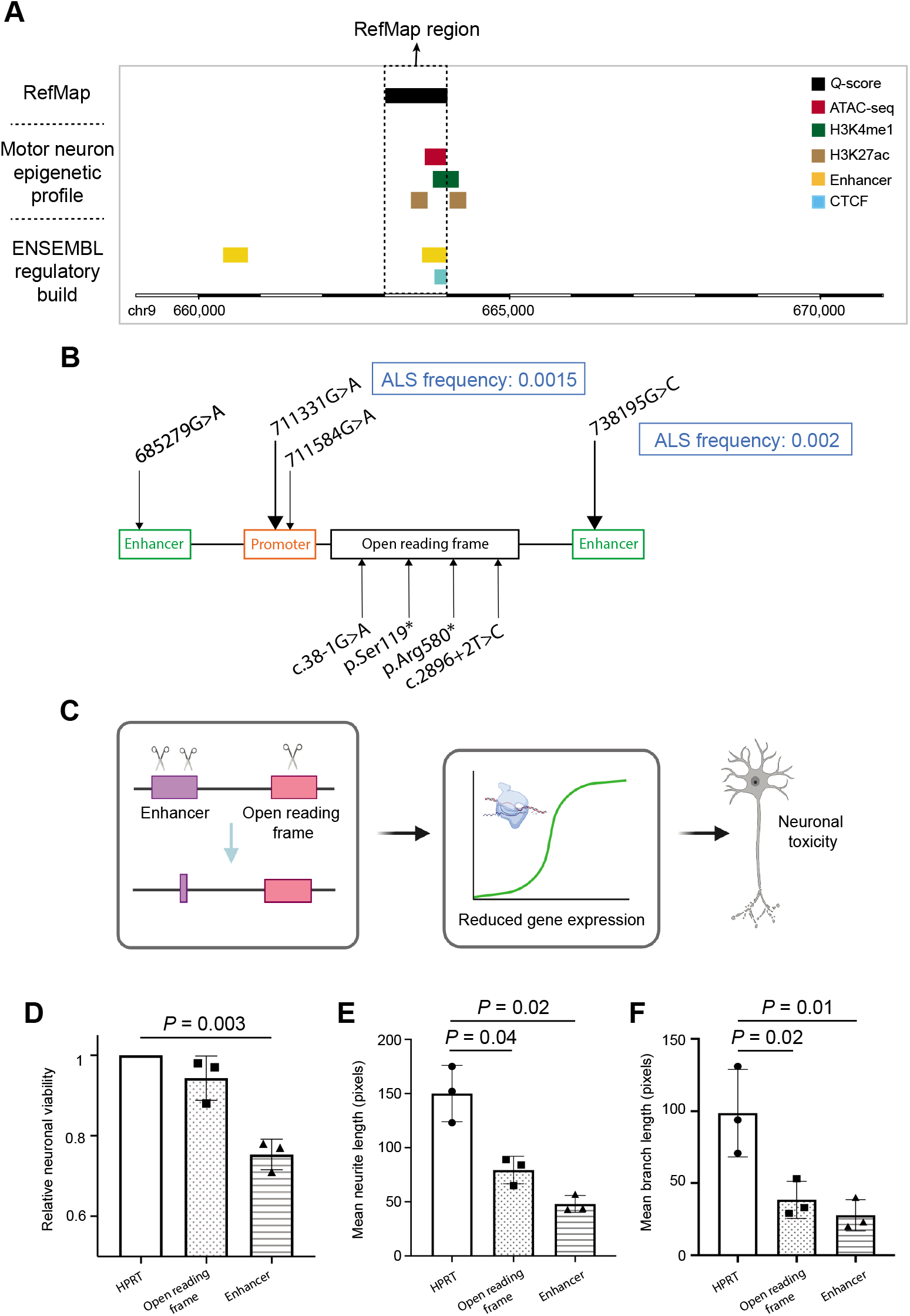

If reduced expression of *KANK1* is linked to MN toxicity, then it is reasonable to expect other loss-of-function (LoF) *KANK1* mutations to be associated with an increased risk of ALS. Thus far RefMap has utilized common genetic variants from a GWAS study (van Rheenen et al., 2016) so, to further investigate *KANK1,* here we performed rare variant burden tests. Rare variant analysis utilized whole-genome sequencing (WGS) data from 5,594 sporadic ALS patients and 2,238 controls (Project MinE ALS Sequencing Consortium, 2018). We filtered for rare, deleterious variants within *KANK1* enhancer, promoter and coding regions based on evolutionary conservation, functional annotations and population frequency (Huang et al., 2017; Karczewski et al., 2020; Rentzsch et al., 2019; Ritchie et al., 2014) (**Methods**). Enhancer and promoter regions for *KANK1* were defined as previously described (Cooper-Knock et al., 2020; Fishilevich et al., 2017). Enhancer and promoter regions were independently enriched with ALS-associated rare deleterious variants (P<0.05, SKAT; **Fig. 5b**) (Lee et al., 2012; Wu et al., 2011), and nonsense coding variants were absent from controls and present in a small number (n=4) of ALS patients. Across all three regions, there was significant enrichment of rare deleterious variants in ALS patients compared to controls (*P*=0.003, Stouffer’s method (Whitlock, 2005); **Fig. 5b**). The observation of both rare and common ALS-associated genetic variation in independent datasets utilizing independent methodology strongly suggests *KANK1* is a new ALS risk gene.

*KANK1* was located within a distinct module (M826, 687 genes; **Supplementary Fig. 3b**) in our network analysis, and this module is enriched with RefMap genes (*P*=5.6e-3, hypergeometric test), but not after multiple testing correction. Functionally the *KANK1*-module is highly expressed in normal MNs (**Fig. 4f**), and is enriched for biological functions centered on the distal axon and synapse (**Supplementary Fig. 3c**), which are consistent with other RefMap-enriched modules.

### Experimental validation of KANK1 in ALS development

To further investigate the role of *KANK1* in ALS, we experimentally determined the effect of ALS-associated genetic variation on gene expression and neuronal health (**Fig. 5c**). We used CRISPR/SpCas9 editing of SH-SY5Y neurons to recapitulate ALS-associated regulatory and coding mutations.

We discovered a high density of ALS-associated genetic variants within a region at chr9:663001-664000, which also contains an independently validated enhancer element (**Fig. 5a**). To replicate disruption of this sequence, we designed gRNAs to target protospacer adjacent motif (PAM) sites up- and downstream so as to delete the entire region (Zheng et al., 2014) (**Methods**). In addition, our rare variant analysis identified ALS-associated nonsense mutations in ALS cases but not in controls, therefore we also targeted a PAM site within *KANK1* exon 2 so as to introduce a series of indels (**Methods**). Sanger sequencing and waveform decomposition analysis(Hsiau et al.) in undifferentiated SH-SY5Y cells confirmed the exon 2 editing efficiency (**Supplementary Figs. 4a** and **4b**) and the deletion of the enhancer sequence (**Supplementary Fig. 4c**). For experimental evaluation, a commercially available control gRNA targeting HPRT served as a negative control. CRISPR/SpCas9-edited SH-SY5Y cells were differentiated to a neuronal phenotype, and successful differentiation was confirmed by altered expression of PAX6 (**Supplementary Fig. 4d)** (Forster et al., 2016) and increased total dendritic length (*P*=0.046, paired Student’s *t*-test; **Supplementary Fig. 4e)** (Forster et al., 2016). Differentiated cells were harvested and RNA was extracted for qPCR. We confirmed the reduced expression of *KANK1* mRNA in both exon and enhancer edited neurons (**Supplementary Fig. 4f**). Furthermore, the reduction in *KANK1* expression was associated with a trend towards reduced neuronal viability in exon edited cells, and with a significant reduction in neuronal viability in enhancer edited cells (exon: *P*=0.1, enhancer: *P*=0.003, paired Student’s *t*-test; **Fig. 5d**). Finally, neurons with reduced expression of *KANK1* exhibited shorter neurites (exon: *P*=0.04, enhancer: *P*=0.02, paired Student’s *t*-test; **Fig. 5e**) with reduced branch length (exon: *P*=0.02, enhancer: *P*=0.01, paired Student’s *t*-test; **Fig. 5f**). In all instances, measures of neuronal toxicity are correlated with *KANK1* expression (**Supplementary Fig. 4f**), which in turn reflects editing efficiency (**Supplementary Figs. 4a-c**). These experimental observations collectively demonstrate the neuronal toxicity focused on the axon caused by ALS-associated genetic variants in *KANK1,* and further support *KANK1* as a new ALS risk gene.

## DISCUSSION

Study of the genetic architectures of complex diseases has been greatly advanced by large GWAS studies. However, many of these studies have not considered cell-type-specific aspects of genomic function, which is particularly relevant for noncoding regulatory sequence (Heinz et al., 2015). This may explain why diseases such as ALS have been linked to relatively few risk genes despite substantial estimates of heritability (Ryan et al., 2019; Trabjerg et al., 2020). Fine-mapping methods have been proposed to disentangle causal SNPs from genetic associations (Benner et al., 2016; Chen et al., 2016b; Hormozdiari et al., 2014; Kichaev et al., 2014; Pickrell, 2014; Schaid et al., 2018), but these approaches are not integrated with cell-type-specific biology (Benner et al., 2016; Hormozdiari et al., 2014), or assume a fixed number of causal SNPs per locus (Chen et al., 2016b; Kichaev et al., 2014; Pickrell, 2014), limiting their power for gene discovery. We have characterized epigenetic features within MNs, which are the key cell type for ALS pathogenesis. Integrating MN epigenetic features with ALS GWAS data in our RefMap model has discovered 690 ALS risk genes, which extends the list of candidate ALS genes by two orders of magnitude. We confirmed the effectiveness of RefMap by direct comparison with three popular fine-mapping methods (Kichaev et al., 2014; Lamparter et al., 2016; de Leeuw et al., 2015), which recovered a maximum of 10 ALS genes. Others have performed more limited epigenetic profiling of motor neurons (Song et al., 2019), but our data are unique with respect to the depth and number of assessments.

Consistent with previous literature, RefMap ALS genes are functionally associated with the distal axon (Frey et al., 2000; Moloney et al., 2014). Several known ALS risk genes are related to axonal function and axonal transport in particular (De Vos and Hafezparast, 2017). Unlike previous literature, our work is based on a comprehensive genome-wide screening and not on a small number of rare variants. As a result, our data suggest that the distal axon may be the site of disease initiation in most ALS patients, and should be the focus of future translational research.

RefMap ALS genes include *KANK1,* which is enriched with common and rare ALS-associated genetic variation across multiple domains and datasets. *KANK1* is functionally related to a number of known ALS genes that are important for cytoskeletal function, including *PFN1, KIF5A* and *TUBA4A.* In particular, *PFN1,* like *KANK1,* is implicated in actin polymerization (Boopathy et al., 2015). Disruption of actin polymerization has been associated with alterations in synaptic organization (Dillon and Goda, 2005), including the neuromuscular junction (NMJ) (Mallik and Kumar, 2018), but also with nucleocytoplasmic transport defects (Giampetruzzi et al., 2019). We have experimentally verified the link between variants identified by RefMap to ALS, and *KANK1* expression. Moreover, we have demonstrated that the reduced expression of *KANK1* in a human CNS-relevant neuron is toxic and produces axonopathy. By contrast, *KANK1* upregulation could be a new therapeutic target for ALS patients with mutations that reduce *KANK1* expression, and possibly more broadly.

In summary, our study provides a general framework that can be applied for the identification of risk genes involved in a large number of complex diseases. With the expansion of genotyping data and increasing understanding of cell-type-specific functions, it should prove valuable to the identification of the genetic underpinnings of many such diseases.

## SUPPLEMENTARY INFORMATION

**Supplementary Figure 1.**
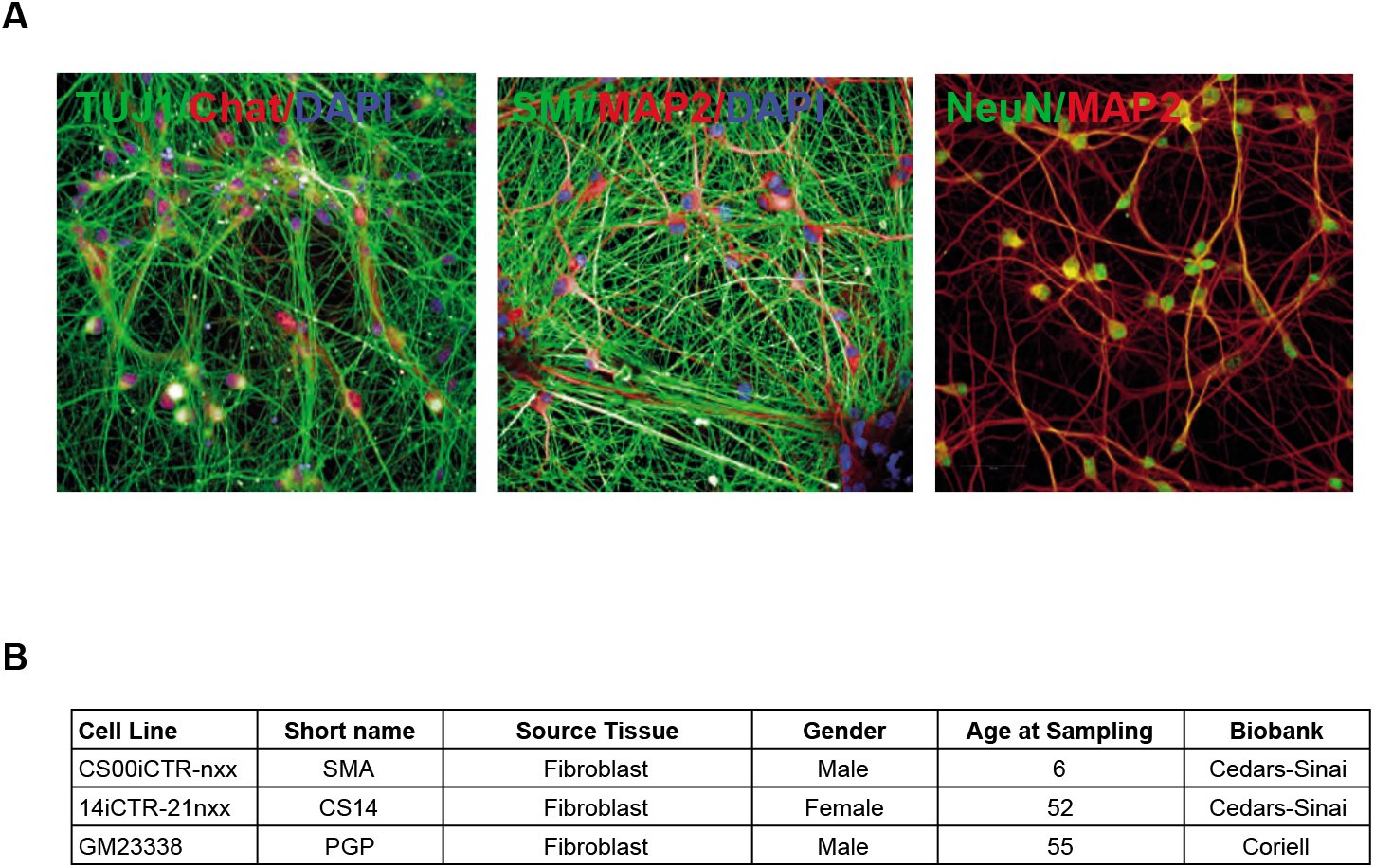
iPSC-derived motor neurons. (**a**) are morphologically consistent with lower motor neurons including expression of appropriate markers. (**b**) iPSC cells were derived from control fibroblasts.

**Supplementary Figure 2.**
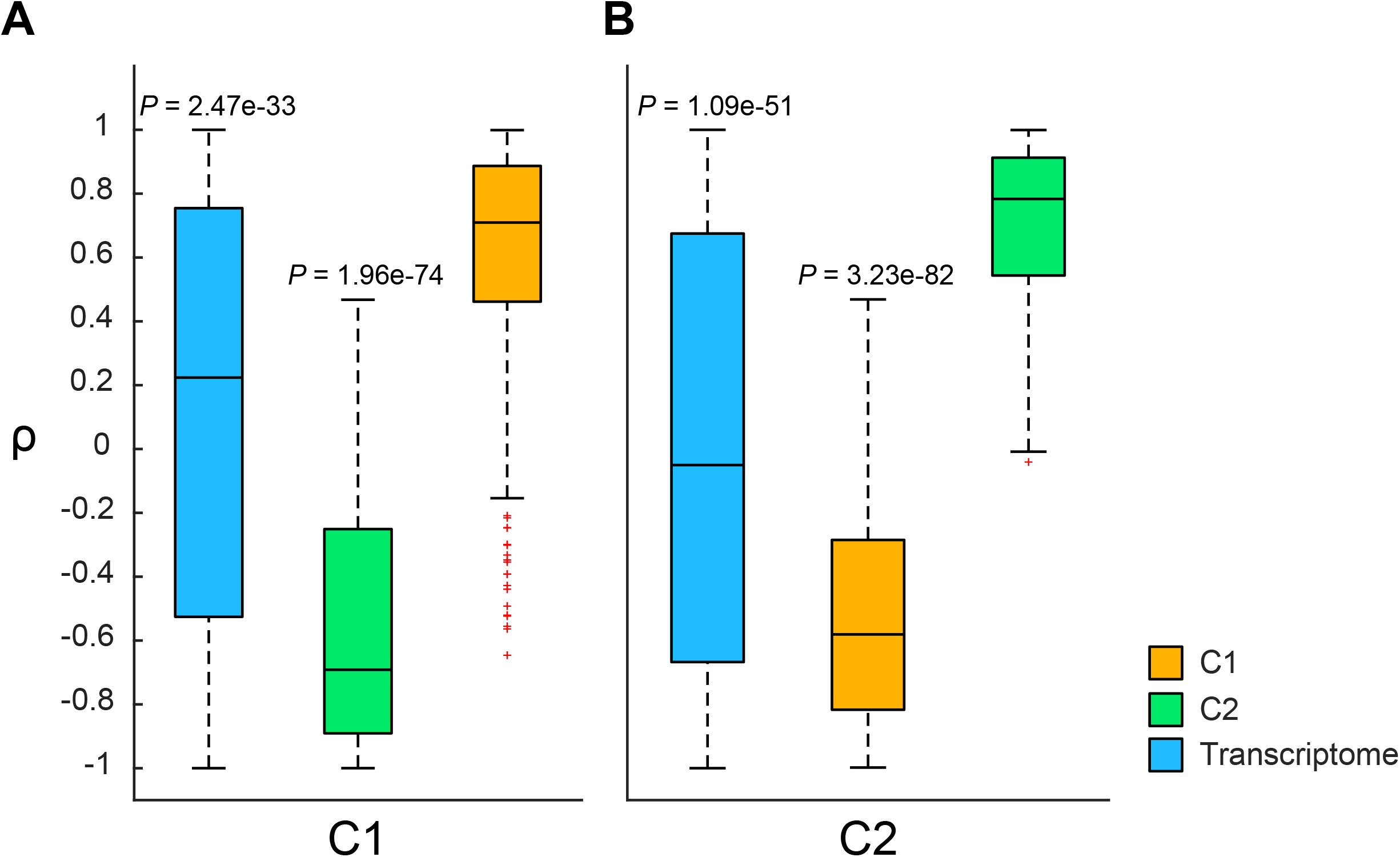
Clustering quality checking for clusters (**a**) C1 and (**b**) C2. The correlations of *Δβ* for individual genes and the mean of the cluster were calculated. The comparisons were performed using the Wilcoxon rank-sum test. Significantly increased between-cluster distance and significantly decreased in-cluster distance were observed, demonstrating the high quality of the clustering.

**Supplementary Figure 3.**
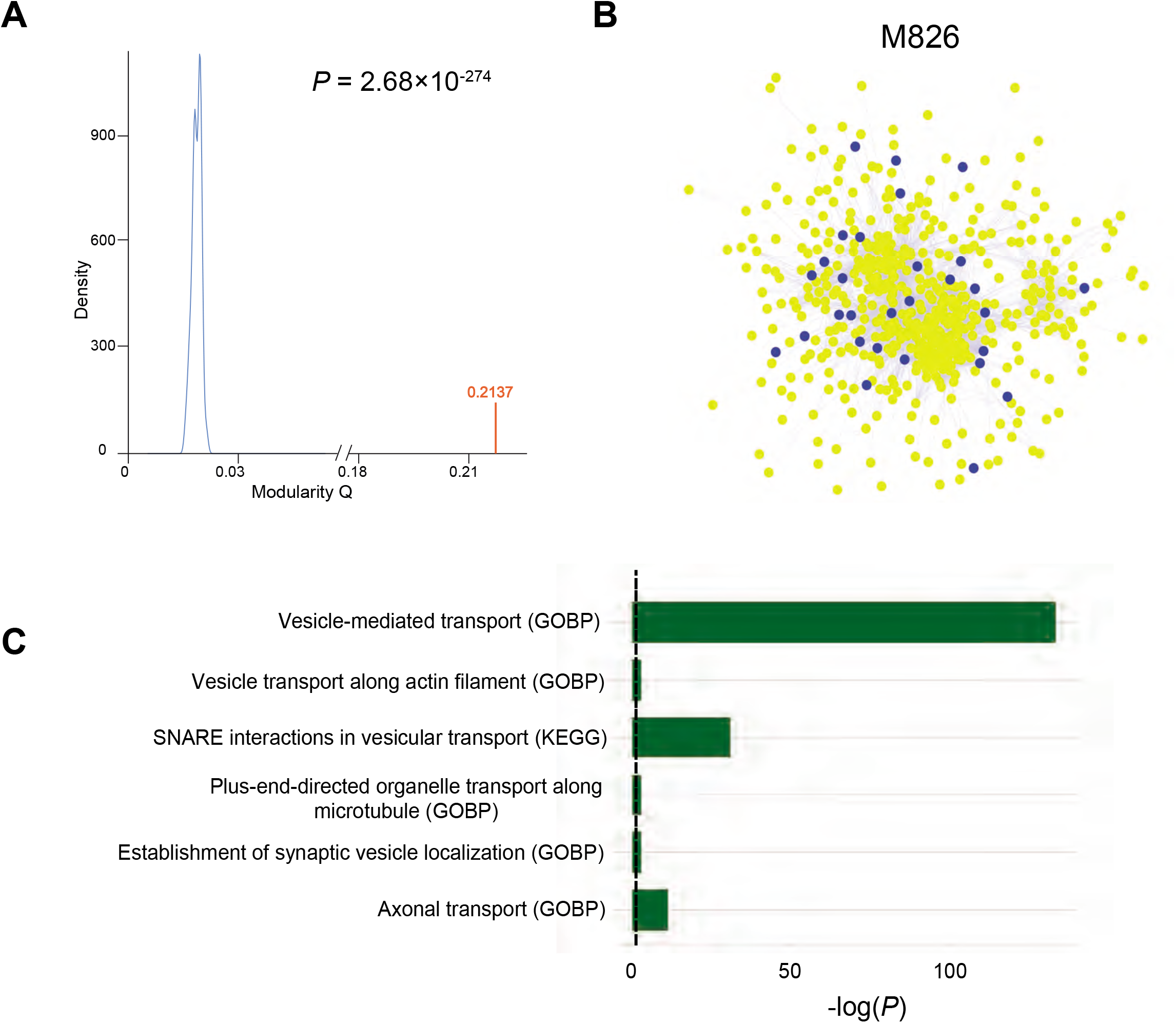
Additional results from network analysis. (**a**) Distribution of modularities after Louvain for the smoothed PPI network (red) and 100 randomized networks (blue). The modularity of our smoothed network is significantly shifted from the randomized network. (**b**) M826 contains *KANK1.* M826 is enriched with RefMap genes (*P*=5.6e-3, hypergeometric test) but not after multiple testing correction. (**c**) M826 is functionally enriched for vesicle transport within the motor neuron axon. GOBP=Gene Ontology Biological Process. Dashed line represents *P*=0.05.

**Supplementary Figure 4.**
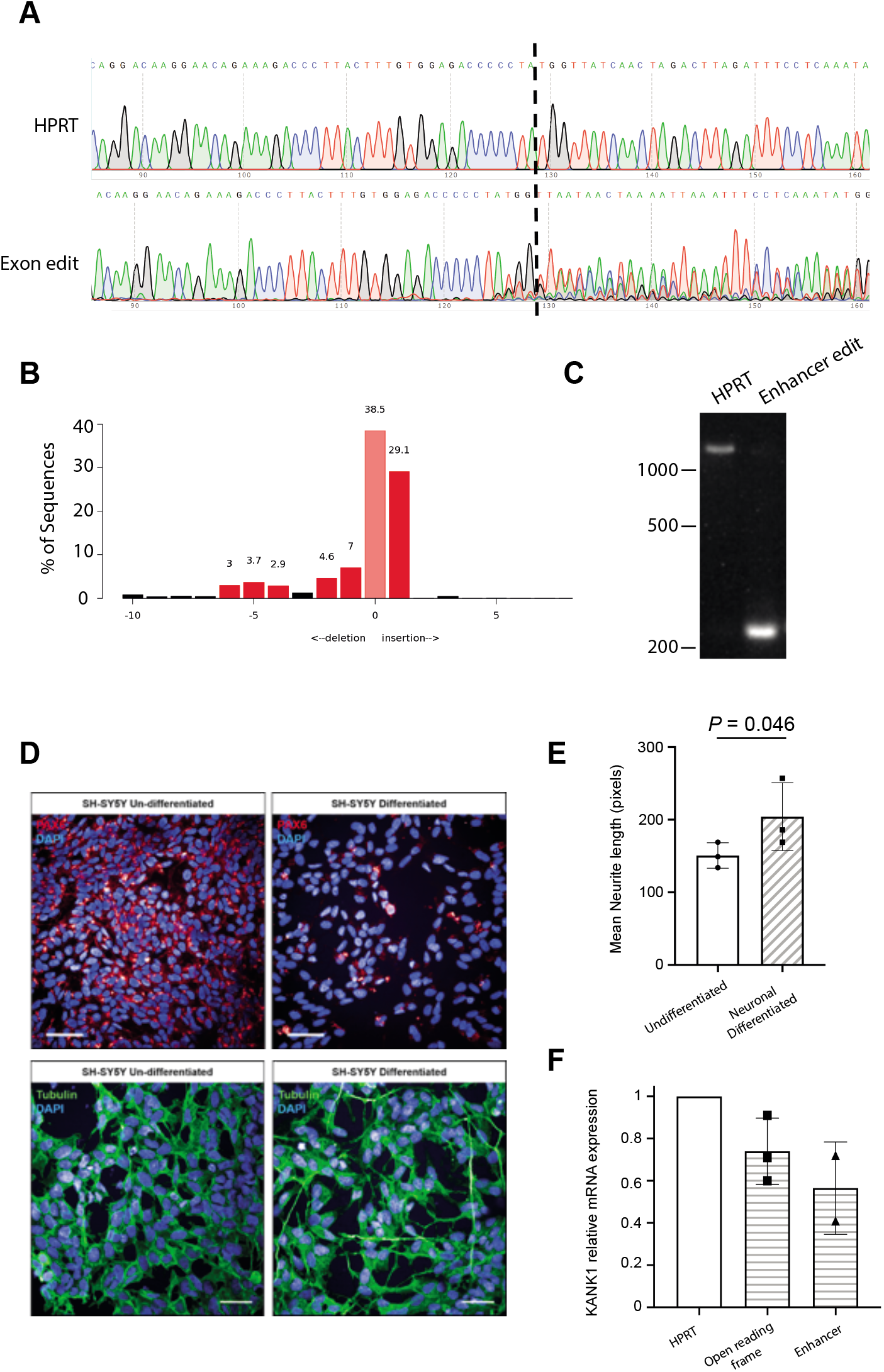
CRISPR-editing of SH-SY5Y cells. (**a**) Sanger sequencing traces demonstrating spCas9 cut site adjacent to PAM and subsequent waveform decomposition in *KANK1* open reading frame edited cells. (**b**) Indel distribution within *KANK1* open reading frame CRISPR-edited SH-SY5Y cells. (**c**) PCR amplification of the relevant genomic segment in enhancer CRISPR-edited SH-SY5Y cells reveals that the chr9:663001-664000 region has been resected compared to HPRT-edited control cells. (**d**) Altered PAX6 expression and (**e**) increased dendrite length confirm the successful neuronal differentiation of SH-SY5Y cells. (**f**) pPCR reveals that the expression of *KANK1* mRNA was reduced in CRISPR-edited SH-SY5Y neurons.

**Supplementary Table 1**

Quality control measures for RNA-seq of iPSC-derived motor neurons

**Supplementary Table 2**

Quality control measures for ATAC-seq of iPSC-derived motor neurons

**Supplementary Table 3**

Quality control measures for histone ChIP-seq of iPSC-derived motor neurons

**Supplementary Table 4**

Quality control measures for Hi-C of iPSC-derived motor neurons

**Supplementary Table 5**

ALS-associated regions identified by RefMap including Q-scores

**Supplementary Table 6**

690 ALS-associated genes identified by RefMap

**Supplementary Table 7**

Manually curated ALS gene list with evidence for association including references

**Supplementary Table 8**

ALS genes discovered by analysis of GWAS data using MAGMA, Pascal and PAINTOR.

**Supplementary Table 9**

Clusters of RefMap homologs in transcriptome data from *SOD1*-G93A ALS mouse model

**Supplementary Table 10**

Smoothed PPI network after preserving top 5% edges predicted by random walk with restart

**Supplementary Table 11**

Gene modules detected from network analysis

**Supplementary Table 12**

Project MinE ALS Sequencing Consortium

**Supplementary Notes**

Mathematical and technical details of RefMap

## ACKNOWLEDGEMENTS

This work used the Genome Sequencing Service Center by Stanford Center for Genomics and Personalized Medicine Sequencing Center, supported by the grant award NIH S10OD025212, and NIH/NIDDK P30DK116074. We acknowledge the Stanford Genetics Bioinformatics Service Center for providing computational infrastructure for this study. We thank W. Rheenen for the explanation of ALS GWAS data. We thank J. Adrian for the help to initiate the project. We also thank J. Zhai and X. Yang for the help with histone ChIP-seq assays, and I. Gabdank and M. Kagda for running the Hi-C pipeline. This project has received funding from the European Research Council (ERC) under the European Union’s Horizon 2020 research and innovation programme (grant agreement n° 772376 - EScORIAL. The collaboration project is co-funded by the PPP Allowance made available by Health~Holland, Top Sector Life Sciences & Health, to stimulate public-private partnerships. This study was also supported by the ALS Foundation Netherlands, by research grants from IWT (n° 140935), the ALS Liga België, the National Lottery of Belgium, the KU Leuven Opening the Future Fund, the National Institutes of Health (CEGS 5P50HG00773504, 1P50HL083800, 1R01HL101388, 1R01-HL122939, S10OD025212, and P30DK116074, UM1HG009442 to M.P.S.), the Wellcome Trust (216596/Z/19/Z to J.C.K.) and NIHR (P.J.S.). We acknowledge support from a Kingsland fellowship (T.M.), the My Name’5 Doddie Foundation (J.F.), and the NIHR Sheffield Biomedical Research Centre for Translational Neuroscience. Biosample collection was supported by the MND Association and the Wellcome Trust (P.J.S.). We are very grateful to those ALS patients and control subjects who generously donated biosamples. We acknowledge transcriptome data provided by the AnswerALS Consortium.

## AUTHOR CONTRIBUTIONS

S.Z., J.C.K. and M.P.S. conceived and designed the study. S.Z. contributed to the design, theoretical analysis and implementation of RefMap. S.Z., J.C.K., A.K.W., M.S., T.M., C.H., H.G.N., J.F., C.S.S., L.F., P.J.S. and M.P.S. were responsible for data acquisition. S.Z., J.C.K., A.K.W., M.S., T.M., C.H., H.G.N., J.F., C.S.S., C.W., J.L., C.E., E.H., L.F., P.J.S. and M.P.S. were responsible for analysis of data. S.Z., J.C.K., A.K.W., M.S., T.M., C.H., H.G.N., J.F., C.S.S., C.W., J.L., C.E., E.H., K.P.K., J.V., L.F., P.J.S. and M.P.S. were responsible for interpretation of data. The Project MinE ALS Sequencing Consortium (**Supplementary Table 12**) was involved in data acquisition and analysis. S.Z., J.C.K. and M.P.S. prepared the manuscript with assistance from all authors. All authors meet the four ICMJE authorship criteria, and were responsible for revising the manuscript, approving the final version for publication, and for accuracy and integrity of the work.

## STAR METHODS

### KEY RESOURCES TABLE

| Reagent or Resource | Source | Identifier |
| --- | --- | --- |
| *Antibodies* |  |  |
| Anti-H3K4me1 | Cell Signaling Technologies | #5326S, lot 3 |
| Anti-H3K4me3 | Cell Signaling Technologies | #9751S, lot 10 |
| Anti-H3K27ac | ActiveMotif | #93133, lot 28518012 |
| α-tubulin | Sigma | #T9026 |
| Anti-Pax6 | Abcam | #ab5790 |
| *Chemicals, Peptides and Recombinant Proteins* |  |  |
| Thiazolyl Blue Tetrazolium Bromide (MTT) | Sigma-Aldrich | #M2128 |
| Alt-R S.p. Cas9 Nuclease V3 | Integrated DNA technologies | #1081059 |
| Alt-R Cas9 Electroporation Enhancer | Integrated DNA technologies | #1075915 |
| Dulbecco's Modified Eagle medium | Lonza | #12-604F |
| KnockOut DMEM/F-12 | ThermoFisher Scientific | #12660012 |
| Neurobasal medium | ThermoFisher Scientific | #12348017 |
| Penicillin-Streptomycin | Sigma | #P4333 |
| Fibronectin | Merck | #FC010 |
| 10x Trypsin | Sigma | #59427C |
| Foetal bovine serum | ThermoFisher Scientific | #10270106 |
| Matrigel | Corning | #356230 |
| mTeSR-Plus Medium | StemCell Technologies | #05825 |
| ReLeSR | StemCell Technologies | #05872 |
| Ethidium bromide solution | Sigma | #E1510 |
| VeriFi mix red | PCRBio | #PB10.42-01 |
| Tri reagent | Sigma | #93289-100ML |
| M-MLV reverse transcriptase | ThermoFisher Scientific | #28025-013 |
| 5x First Strand buffer | ThermoFisher Scientific | #18057-018 |
| 0.1M Dithiothreitol | ThermoFisher Scientific | #707265ML |
| dNTP Mix | ThermoFisher Scientific | #10534823 |
| SYBR Green Brilliant III master mix | Agilent | #600882 |
| Random hexamer primer | ThermoFisher Scientific | #SO142 |
| Purmorphamine | Tocris Bioscience | #4551 |
| StemPro Accutase Cell Dissociation Reagent | Gibco | #A1110501 |
| ROCK inhibitor (Y-27632 dihydrochloride) | Tocris Bioscience | #1254 |
| Compound E | Tocris Bioscience | #6476 |
| NEBNext 2xMasterMix | New England Biolabs | M0541 |
| EDTA | Sigma | #E5134 |
| HEPES | Sigma | #H3375 |
| PMSF protease inhibitor | ThermoFisher Scientific | #36978 |
| Protease inhibitor tablet | Roche | #1697498 |
| Gibco GlutaMAX Supplement | ThermoFisher Scientific | #35050061 |
| TracrRNA | Integrated DNA technologies | #1072533 |
| TE Buffer, RNase-free pH 8 | ThermoFisher Scientific | #AM9849 |
| Dulbecco's Phosphate Buffered Saline | Sigma | #D8537-500ML |
| Triton X-100 | Sigma-Aldrich | #T8787 |
| Normal horse serum | Vector | #S-2000-20 |
| Hoechst 33342 | ThermoFisher Scientific | #62249 |
| All-trans retinoic acid | Sigma | #R2625 |
| BDNF | PeproTech | #450-02 |
| IGF | ThermoFisher Scientific | #PHG0078 |
| CNTF | ThermoFisher Scientific | #PHC7015 |
| N-2 supplement | ThermoFisher Scientific | #17502048 |
| B-27 supplement | ThermoFisher Scientific | #17504001 |
| DMH-1 | Tocris Bioscience | #4126 |
| SB431542 | Tocris Bioscience | #1614 |
| CHIR99021 | Tocris Bioscience | #4423 |
| *Critical Commercial Assays* |  |  |
| Pierce BCA Assay Protein Assay Kit | ThermoFisher Scientific | #23225 |
| GenElute Mammalian Genomic DNA Miniprep Kit | Sigma | #G1N350 |
| Direct-zol RNA Miniprep Kit | Zymo Research | #R2050 |
| Neon Transfection System 10 μL Kit | ThermoFisher Scientific | #MPK1096 |
| Alt-R CRISPR-Cas9 Control Kit, Human, 2 nmol | Integrated DNA technologies | #1072554 |
| QIAquick PCR Purification kit | Qiagen | #28104 |
| MiElute kit. | Qiagen | #28004 |
| KAPA Library Quantification kit | Roche | #07960140001 |
| KAPA HiFi HotStart ReadyMix | Roche | #07958927001 |
| KAPA Library Amplification Primer Mix | Roche | #07958978001 |
| QIAquick Gel Extraction Kit | Qiagen | #28506 |
| Ribo-Zero rRNA depletion kit | Illumina | #20040526 |
| NEBext Ultra RNA prep kit | New England Biolabs | #E7530 |
| *Experimental Models: Cell Lines* |  |  |
| SH-SY5Y | ATCC | Cat.#CRL-2266 |
| *Software and Algorithms* |  |  |
| Sickle v1.200 | https://github.com/najoshi/sickle |  |
| Cutadapt v1.2.1 | https://pypi.org/project/cutadapt/1.2.1/ |  |
| Kallisto v0.46.0 | https://pachterlab.github.io/kallisto/ |  |
| SKAT | https://cran.r-project.org/web/packages/SKAT/index.html |  |
| R v4.0.1 | https://cran.r-project.org/mirrors.html |  |
| snpStats | https://www.bioconductor.org/packages/release/bioc/html/snpStats.html |  |
| VariantAnnotation | https://www.bioconductor.org/packages/release/bioc/html/VariantAnnotation.html |  |
| PLINK V1.90 | http://zzz.bwh.harvard.edu/plink/download.shtml |  |
| PRISM 7 | GraphPad |  |
| ICE CRISPR analysis tool | https://ice.synthego.com/#/ |  |
| CRISPOR guide RNA design tool | http://crispor.tefor.net/ |  |
| CFX Maestro | Bio-Rad |  |
| Harmony Imaging Analysis Software | PerkinElmer |  |
| FIJI (FIJI Is Just ImageJ) | NIH |  |
| MATLAB R2018b | MathWorks |  |
| MAGMA v1.08 | https://ctg.cncr.nl/software/magma |  |
| Pascal | https://www2.unil.ch/cbg/index.php?title=Pascal |  |
| PAINTOR v3.0 | https://github.com/gkichaev/PAINTOR_V3.0 |  |

### RESOURCE AVAILABILITY

#### Lead Contact

Further information and requests for resources and reagents should be directed to and will be fulfilled by the corresponding author, Micheal P. Snyder.

#### Materials Availability

All unique/stable reagents generated in this study are available from the Lead Contact without restriction.

#### Data and Code Availability

Epigenetic profiling of iPSC-derived motor neurons is available at encodeproject.org with the following accession numbers: ENCSR065CER, ENCSR410DWV, ENCSR812ZKP, ENCSR634WYX, ENCSR459PVP, ENCSR913OWV, ENCSR704VZY, ENCSR131HOY, ENCSR516YAD, ENCSR709QRD ENCSR754DRC, ENCSR672RKZ, ENCSR571HAY, ENCSR503HWR, ENCSR207VLY, ENCSR962OTG, ENCSR745TRI, ENCSR595HWK, ENCSR312HLG, ENCSR682BFG, ENCSR680IWU, ENCSR564EFE, ENCSR358AOC, ENCSR698HPK, ENCSR778FKK, ENCSR425FUS, ENCSR489LNU and ENCSR540KQC.

Code is available on request.

### EXPERIMENTAL MODEL AND SUBJECT DETAILS

#### Study cohorts

iPSC-cells were derived from fibroblasts obtained from three neurologically normal controls of different ages: 55-year old male, a 52-year old female and a 6-year old male (**Supplementary Fig. 1b**). GWAS summary statistics were previously published (van Rheenen et al., 2016). The 6,180 patients and 2,370 controls included in this study were recruited at specialized neuromuscular centers in the UK, Belgium, Germany, Ireland, Italy, Spain, Turkey, the United States and the Netherlands (Project MinE ALS Sequencing Consortium, 2018). Patients were diagnosed with possible, probable or definite ALS according to the 1994 El-Escorial criteria (Brooks, 1994). All controls were free of neuromuscular diseases and matched for age, sex and geographical location.

The study was approved by the South Sheffield Research Ethics Committee. Also, this study followed study protocols approved by Medical Ethical Committees for each of the participating institutions. Written informed consent was obtained from all participating individuals. All methods were performed in accordance with relevant national and international guidelines and regulations.

#### SH-SY5Y neuroblastoma cells

Human SH-SY5Y neuroblastoma cells were cultured in Dulbecco’s Modified Eagle’s Medium (DMEM) (Lonza) supplemented with 10% (v/v) foetal bovine serum (FBS) (Thermo-Fisher Scientific), 50 units/mL of penicillin and 50 μg/mL of streptomycin. Cell lines were maintained at 5% CO2 in a 37°C incubator and split every 3-4 days. All experimental work was performed using cells within the range of 7-32 passages.

### METHOD DETAILS

#### Cell culture

Human induced pluripotent stem cells iPSCs were maintained in Matrigel-coated plates (Corning) according to the manufacturer’s recommendations in complete mTeSR-Plus Medium (StemCell Technologies). The culture medium was replaced daily and confirmed mycoplasma free. Cells were passaged every four to six days as clumps using ReLeSR an enzyme-free reagent for dissociation (StemCell Technologies) according to the manufacturer’s recommendations. For all the experiments in this study, iPSCs were between passage 20 and 32.

#### iPSC-derived motor neuron differentiation

iPSCs derived from unaffected controls were differentiated to motor neurons using the modified version of the dual SMAD inhibition protocol (Du et al., 2015). Briefly iPCS cells were transferred for Matrigel-coated plate (Corning). On the day after plating (day 1), after the cells had reached ~100% confluence, the cells were washed once with PBS and then the medium was replaced for neural medium (50% of KnockOut DMEM/F-12, 50% of Neurobasal), 0.5× N2 supplement (ThermoFisher), 1x Gibco GlutaMAX Supplement (ThermoFisher), 0.5x B-27 (ThermoFisher), 50 U ml^−1^ penicillin and 50 mg ml^−1^ streptomycin, supplemented with SMAD inhibitors (DMH-1 2μM; SB431542 10 μM; and CHIR99021 3 μM).

The medium was changed every day for 6 days, on day 7, the medium was replaced for neural medium supplemented with DMH-1 2 μM, SB431542-10 μM and CHIR 1 μM, All-Trans Retinoic Acid 0.1 μM (RA), and Purmorphamine 0.5 μM (PMN), the cells were kept in this medium until day 12 when is possible to see a uniform neuroepithelial sheet, the cells were split 1:6 with Accutase (Gibco), onto matrigel substrate in the presence of 10 μM of ROCK inhibitor (Y-27632 dihydrochloride, Tocris), giving rise to a sheet of neural progenitor cells (NPC). After 24 hours of incubation the medium was changed for neural medium supplemented with RA 0.5 μM and PMN 0.1 μM, the medium was changed every day for more 6 days. On day 19 the motor neuron progenitors were split with accutase onto to matrigel-coated plates and the medium was replaced for neural medium supplemented with RA 0.5 μM, PMN 0.1 μM, compound E 0.1 μM (Cpd E, Tocris), BDNF 10ng/mL, CNTF 10ng/mL and IGF 10ng/mL until day 28. On day 29, the media was replaced for Neuronal media (Neurobasal media supplemented with 1% of B27, BDNF 10ng/mL, CNTF 10ng/mL and IGF 10ng/mL). The cells were then fed alternate days with neuronal medium until day 40.

#### ATAC-seq

50,000 viable motor neurons were spun down at 500 RCF at 4°C for 5 min. Supernatant was discarded. 50 μl cold ATAC Resuspension Buffer (RSB) (10 mM Tris-HCl pH 7.4, 10 mM NaCl, 3 mM MgCl2, sterile H2O) containing 0.1% NP40, 0.1% Tween-20, and 0.01% Digitonin was added and carefully mixed. Tubes were incubated on ice for 3 min. 1 ml of cold ATAC-RSB containing 0.1% Tween-20 was added and the tubes were inverted three times. Nuclei were spun down at 500 RCF for 10 min at 4°C. Supernatant was aspirated. Cell pellet was resuspended in 50 μl of transposition mix (25 μl 2x TD buffer, 2.5μl transposase (100 nM final), 16.5 μl PBS, 0.5 μl 1% digitonin, 0.5 μl 10% Tween-20, 5 μl H2O) by pipetting up and down 6 times. TD buffer consists of 20 mM Tris-HCl pH 7.6, 10 mM MgCl2, 20% DMF, sterile H2O. pH was adjusted with acetic acid before adding DMF. The reaction was incubated at 37°C for 30 minutes in a thermomixer while shaking at 1000 RPM. Reaction was cleaned up with a Qiagen MiElute kit. DNA was eluted in 20 μL elution buffer. DNA was amplified using the NEBNext 2xMasterMix. Cycling conditions: 5 min at 72°C, 30 sec at 98°C, followed by 5 cycles of 10 sec at 98°C, 30 sec at 63°C and 1 min at 72°C, hold at 4°C. 5μl (10% of the pre-amplified mixture) were used for qPCR to determine the number of additional cycles needed (3.76 μL H2O, 0.5 μL 25 μM Primer1, 0.5 μL 25 μM Primer2, 0.24 μL 25x SYBR Green, 5 μL NEBNext MasterMix). Cycling conditions: 30 sec at 98°C, followed by 20 cycles of 10 sec at 98°C, 30 sec at 63°C and 1 min at 72°C, hold at 4°C. Amplification profiles were assessed as previously described (Buenrostro et al., 2015). The remainder of the pre-amplified DNA (45μL) was used to run the required number of additional cycles. The final PCR reaction was cleaned up using Qiagen MinElute kit and eluted in 20 μl H2O. Libraries were quantified with the KAPA Library Quantification kit (Roche) and sequenced on a NovaSeq 6000 system (Illumina). Raw data were processed with the ENCODE 4 pipeline for ATAC-seq according to ENCODE 4 standards (https://www.encodeproject.org/atac-seq/). All samples exceeded ENCODE 4 standards for % mapped reads, enrichment of transcription start sites, the fraction of reads that fall within peak regions (FRiP), and reproducibility between technical replicates (**Supplementary Table 1**).

Files are available at encodeproject.org with the following accession numbers: ENCSR065CER, ENCSR410DWV, ENCSR812ZKP, ENCSR634WYX, ENCSR459PVP, ENCSR913OWV, ENCSR704VZY, ENCSR131HOY, ENCSR516YAD, ENCSR709QRD.

#### Histone ChIP-seq

5 million motor neurons were crosslinked and resuspended in 10 mL of cold L1 buffer (50mM Hepes KOH, pH 7.5, 140mM NaCl, 1mM EDTA, 10% Glycerol, 0.5% NP-40, 0.25% Triton X-100, dH_2_O, 1 protease inhibitor tablet (Roche) per 50ml buffer). Cells were incubated on a rocking platform at 4°C for 10 minutes and spun down at 3000 rpm at 4°C for 10 minutes. Pellets were resuspended in 10 mL of L2 buffer (200mM NaCl 1mM EDTA pH 8 0.5mM EGTA 10mM Tris, pH 8, dH_2_O, 1 protease inhibitor tablet (Roche) per 50ml buffer, room temperature). Tubes were incubated at room temperature for 10 minutes and spun down at 3000 rpm for 10 minutes at 4°C. Nuclei were resuspended in 3 mL 1X RIPA buffer and incubated on ice for 30 minutes. Samples were sonicated with Branson 250 Sonifier to shear the chromatin. 3 mL of sheared chromatin lysate were transferred to two 2 mL tubes and spun down at 14,000 rpm at 4°C for 15 minutes. 50 μL were saved from each replicate and pooled as input (no antibody added, kept at −20°C). 2 μL histone modification antibody was added to each 3 mL lysates and incubated at 4°C on a neutator for 12-16 hours. The following antibodies were used: H3K4me1 (Cell Signaling Technologies), H3K4me3 (Cell signaling technologies), H3K27ac (ActiveMotif). 80 μL of Protein A/G-agarose for each sample were washed twice with 1 mL of ice cold 1X RIPA buffer, spun down at 5000 rpm for 1 minute at 4°C and resuspended in 80μL in 1x RIPA buffer. Beads were added to tubes containing Ag-Ab complex (80 μ L 1X RIPA to wash out the beads) and incubated for 1 hour at 4°C with neutator rocking. Tubes were spun down at 1500 rpm for 3 minutes, beads were washed 3 times 15 minutes each with 10 mL of fresh, ice cold 1x RIPA buffer supplemented per 50 mL with 1 protease inhibitor tablet, 250 μL of 100 mM PMSF, 50 μL of 1M DTT, 2 ml of phosphatase inhibitor (sodium pyrophosphate 1mM, sodium orthovanadate 2mM, sodium fluoride 10mM). Afterwards, beads were washed once with ice cold 1 x PBS for 15 minutes. Beads were resuspended in 1200 μL ice cold 1x PBS, transferred to an 1.5mL Eppendorf tube and spun down at 5000 rpm for 1 minute. PBS was removed and 100 μL of Elute 1 solution (1% SDS, 1x TE, dH_2_O) was added to resuspend beads and tubes were incubated at 65°C for 10 minutes with gentle mixing every 2 minutes. Beads were spun down at 5000 rpm for 1 minute at room temperature and the supernatant was kept as Elute 1. 150 μL of Elute 2 solution (0.67% SDS, 1x TE) was added to the bead pellets and incubated at 65°C for 10 minutes with gentle vortexing. After spinning down for 1 minute at 5000 rpm, the second elute was combined with the first. Input DNA was thawed and 150 μL of Elute 1 solution was added. All samples incubated at 65°C overnight to reverse cross-linking. 250 μL 1X TE containing 100 μg RNase was added to each sample and incubated for 30 minutes at 37°C. 5 μL of 20 mg/mL Proteinase K was added to each sample and incubated at 45°C for 30 minutes. After transferring samples to 15 mL tubes, DNA was purified (Qiaquick PCR purification kit, Qiagen). DNA was eluted in elution buffer (50μL for input, 35μL for ChIP sample).

The following components were combined and mixed in a microfuge tube: ChIP DNA to be end-repaired (25ng) 34 μL, 5 μL 10X End-Repair Buffer, 5 μL 2.5 mM dNTP Mix, 5 μL10 mM ATP, 1 μL End-Repair Enzyme Mix. The mixture was incubated at room temperature for 45 minutes. DNA was purified (MinElute PCR purification kit, Quiagen) and eluted in 19 μL EB. Adapter ligated DNA was run on a 2% EX-Gel and excised in the range of 450-650 bp with a clean scalpel. DNA was purified (Gel extraction kit, Quiagen) and eluted in 20 μL EB. The following components were mixed in a PCR tube: 20 μL of purified DNA, 25 μL KAPA HiFi HotStart ReadyMix (2X), 5 μL KAPA Library Amplification Primer Mix (10X). DNA was amplified with the following conditions: 45 sec at 98°C, 15x [15 sec at 98°C, 30 sec at 60°C, 30 sec at 72°C], 60 sec at 72°C, hold at 4°C. The PCR product was purified (MinElute PCR purification kit, Quiagen) and eluted in 19 μL EB. DNA was run on a 2% EX-Gel and excised in the range of 300-450 bp (or brightest smear) with a clean scalpel. DNA was purified (Qiaquick Gel extraction kit, Quiagen) and eluted in 12 μL EB. Library concentration was measured using Qubit and each library was run on the Bioanalyzer. Equal concentrations of different barcoded libraries were pooled and sequenced on a NovaSeq 6000 system (Illumina). Raw data were processed with the ENCODE 4 pipeline for Histone ChIP-seq according to ENCODE 4 standards (https://www.encodeproject.org/chip-seq/histone/). All samples exceeded ENCODE standards for % mapped reads, the fraction of reads that fall within peak regions (FRiP), and reproducibility between technical replicates (**Supplementary Table 2**)

Files are available at encodeproject.org with the following accession numbers: ENCSR754DRC, ENCSR672RKZ, ENCSR571HAY, ENCSR503HWR, ENCSR207VLY, ENCSR962OTG, ENCSR745TRI, ENCSR595HWK, ENCSR312HLG, ENCSR682BFG, ENCSR680IWU, ENCSR564EFE, ENCSR358AOC, ENCSR698HPK, ENCSR778FKK, ENCSR425FUS, ENCSR489LNU, ENCSR540KQC

#### Hi-C

We generated Hi-C libraries following the protocol previously described(Rao et al., 2014, 2017). In brief, 2-5 million cells were crosslinked with formaldehyde. Nuclei were permeabilized and DNA was digested with 100U of MboI. DNA fragments were labelled with biotinylated nucleotides. Ligated DNA was purified and sheared to a length of 300-500 bp after reverse cross-linking. Ligation junctions were pulled-down with magnetic streptavidin beads. Libraries were amplified by PCR and purified. Library concentrations were measured (Qubit). Hi-C libraries were paired-end sequenced on a NovaSeq 6000 system (Illumina). Raw data were processed with the ENCODE 4 pipeline for Hi-C according to ENCODE 4 standards (https://www.encodeproject.org/documents/75926e4b-77aa-4959-8ca7-87efcba39d79/).

#### RNA-seq

RNA libraries were prepared by first depleting ribosomal RNA using the Illumina Ribo-Zero rRNA depletion kit. Strand-specific libraries were then prepared using NEBext Ultra RNA prep kit. RNAseq libraries were paired-end sequences on a NovaSeq 6000 system (Illumina). Minimum 80 million reads were obtained per sample. The raw Fastq files were trimmed for the presence of Illumina adapter sequences using Cutadapt v1.2.1 (Martin, 2011). The reads were further trimmed using Sickle v1.200 with a minimum window quality score of 20. Reads shorter than 15 bp after trimming were removed. If only one of a read pair passed this filter, it was included in the R0 file. Reads were aligned to hg19 transcripts (n=180,253) using Kallisto v0.46.0 (Bray et al., 2016).

#### CRISPR/Cas9 editing of SH-SY5Y cells

Guide RNAs (gRNAs) were designed using the Crispor tool to target *KANK1* regulatory and coding regions. Design was guided by proximity to patient enhancer mutation sites, available protospacer adjacent motifs (PAM), and predicted on- and off-target efficiencies. gRNAs targeting within 30bp either side of the patient enhancer mutation site (chr9:663,001-664,000, hg19) were considered and screened for editing efficiency. One pair of guide sequences (5’ -UCAUGGGAACUCUUCAAAUA-3’ and 5’- UCAUGGGAACUCUUCAAAUA-3’) was most efficient and chosen for subsequent experimentation. Validated, commercially available CRISPR control targeting HPRT (IDT) and *KANK1* exon-targeting (ThermoFisher Scientific, 5’- GUCUAGUUGAUAACCAUAGG-3’) gRNAs were also obtained. gRNA duplexes were assembled from tracrRNA and crRNA in a thermocycler according to manufacturer’s instructions under RNAse-free conditions. Cells were cultured to ensure 70-90% confluency on the day of transfection. 1ml antibiotic-free DMEM (Lonza) was prepared and incubated in 24-well plates at 37°C. CRISPR/Cas9 Ribonucleoproteins were formed by complexing 240ng gRNA duplex with 1250ng Alt-R V3 Cas9 Protein (IDT) in 10μL buffer R (from 10μL Neon transfection kit, ThermoFisher Scientific) - a 1:1 molar ratio - for 10 minutes. 100,000 viable cells were aliquoted per transfection and centrifuged at 400 x g for 4 minutes. Cells were washed in calcium- and magnesium-free Dulbecco’s Phosphate Buffered Saline (Sigma) and centrifuged at 400 x g for 4 minutes. Cell pellets were resuspended in 10μL buffer R containing Cas9 protein and gRNA duplexes. 2μL of 10.8μM electroporation enhancer (IDT) was added and the solution mixed thoroughly to ensure a suspension of single cells. 10μL of this mixture was loaded into a Neon transfection system (ThermoFisher Scientific) and electroporated according to manufacturer’s instructions (1200V, 3 pulse, 20s pulse width for SH-SY5Y cells). Cells were then transferred to pre-warmed media in 24-well plates.

#### Determining CRISPR editing efficiency

Genomic DNA was isolated from CRISPR-edited and control cells using a GenElute Mammalian DNA Miniprep Kit (Sigma) according to manufacturer’s instructions. A ~400bp region around the expected cas9 cut site was amplified by polymerase chain reaction using VeriFi mix (PCRbio). Expected amplification was confirmed using gel electrophoresis, and the products were Sanger-sequenced. Sequencing trace files were uploaded to ICE (https://ice.synthego.com) and an indel efficiency calculated.

#### Quantitative PCR (RT-PCR)

Cells were cultured until at least 70% confluent, lysed on ice using an appropriate volume of Tri Reagent (Sigma) for 5 minutes and transferred to 1.5ml RNAse-free tubes. Total RNA was extracted using a Direct-zol RNA Miniprep Kit (Zymo) according to manufacturer’s instructions, and RNA concentration confirmed using a NanoDrop spectrophotometer (ThermoFisher Scientific). 2μg of total RNA was then converted to cDNA by adding 1μL 10mM dNTPs, 1μL 40μM random hexamer primer (ThermoFisher Scientific), and DNAse/RNAse-free water to a total volume of 14μL. This mixture was heated for 5 minutes at 70°C then placed on ice for 5 minutes. 4μL of 5x FS buffer, 2μL 0.1M DTT, and 1μL M-MLV reverse transcriptase (ThermoFisher Scientific) were then added and cDNA conversion performed in a PCR thermocycler (37°C for 50 minutes, 70°C for 10 minutes). cDNA was amplified using RT-PCR with Brilliant III SYBR Green (Agilent) as per manufacturer’s instructions. Ct analysis was performed using CFX Maestro software (BioRad). GAPDH was chosen as a reference gene because expression is relatively stable in SH-SY5Y cells (Hoerndli et al., 2004).

#### SH-SY5Y neuronal differentiation

Human SH-SY5Y neuroblastoma cells were seeded at densities of either 5×10^4^ cells per well of a 6-well culture plate, or 2×10^3^ cells per well of a 96-well culture plate in DMEM (Lonza) supplemented with 10% (v/v) FBS, 50 units/mL penicillin and 50 μg/mL of streptomycin. 24 hours after seeding the media was changed to DMEM supplemented with 5% (v/v) FBS, 50 units/mL penicillin, 50 μg/mL of streptomycin, 4mM l-glutamine and 10μM retinoic acid. After 72 hours, the medium was switched to neurobasal media (ThermoFisher Scientific) containing 1% (v/v) N-2 supplement 100x, 50 units/mL penicillin, 50 μg/mL of streptomycin, 1% l-glutamine and 50ng/mL human BDNF. Cells were cultured for an additional 3 days until fully differentiated.

#### Immunocytochemistry

SH-SY5Y cells were fixed with 4% paraformaldehyde for 15 minutes and washed 3x with PBS. Cells were blocked in 5% normal horse serum containing 0.1% Triton X-100 for 1 hour at RT. All primary antibodies were diluted in blocking solution (α-tubulin, 1:2000; anti-Pax6, 1:200). Cells were incubated in the primary antibody for 2 hours at RT and washed 3x in PBS before incubation in the appropriate secondary antibody (1:1000 in PBS) for 1 hour at RT. Nuclear counterstain (Hoechst 33342) was applied for 10 minutes followed by a 3x wash in PBS. Cells were imaged using an Opera Phenix High Content Screening System (PerkinElmer).

#### MTT assays

A colorimetric assay using 3-(4, 5-dimethylthiazol-2-yl)-2, 5-diphenyltetrazolium bromide (MTT) dye was used to assess neuronally differentiated SH-SY5Y cellular metabolic activity and hence neuronal viability. 55 μL of 5mg/mL of MTT reagent in PBS was added per well of a 24-well culture plate and incubated at 37°C for 1 hour. 550 μL of unprecipitated 20% SDS in 50% di-methyl formamide (DMF) + dH_2_O (pH 7.4) was added per well and mixed thoroughly to lyse the cells. Cells were incubated in a dark environment on an orbital shaker for 1 hour. The colorimetric change was measured using a PHERAstar FS spectrophotometer (BMG Biotech), and absorbance readings taken at 590nm were normalized to media-only wells. Mean absorbance readings were calculated for each biological repeat and expressed as a percentage of controls.

### QUANTIFICATION AND STATISTICAL ANALYSIS

#### Model design and inference of RefMap

In this study, allele *Z*-scores were calculated as *Z=b/se,* where *b* and *se* are effect size and standard error, respectively, and they were estimated from a mixed linear model in the ALS GWAS study (Nicolas et al., 2018; van Rheenen et al., 2016). Given allele *Z*-scores and the epigenetic profiling of iPSC-derived motor neurons, we were interested in predicting causal associations of individual genomic regions with ALS risk. Suppose we have *K* 1Mb LD blocks with non-zero alleles, whose approximate between-block independence has been verified in previous literature(Loh et al., 2015). Also suppose each LD block contains *J_k_* (*k*=1,…, *K*) 1kb regions and each region harbors *I_j,k_* (*j*=1,…, *J_k_, I_j,k_*>0) SNPs. We further denote the *Z*-score for the *i*-th SNP in the *j*-th region of the *k*-th block as *Z_i,j,k_* (*i*=1,…, *I_j,k_*). Under a linearity hypothesis, we can prove that ***Z**_k_* follows a multivariate normal distribution (**Supplementary Notes**), i.e.,

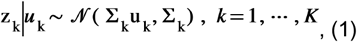

in which ***u**_k_* are the effect sizes of individual SNPs that can be expressed as

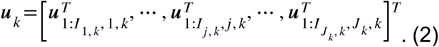

Moreover, in Eq. (1) 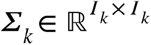 represents the in-sample LD matrix comprising of the pairwise Pearson correlation coefficients between SNPs within the *k*-th block, where *I_k_* is the total number of SNPs given by 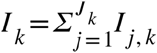. Here, since we have no access to the individual genotypes, we used EUR samples from the 1000 Genomes Project to estimate *∑_k_* (i.e., out-sample LD matrix).

Here, the latent variables ***u**_k_* can be treated as the disentangled *Z*-scores from LD confounding, leaving the right place for independence assumption and facilitating downstream modelling. Indeed, we assume *U_i,j,k_* (*i*=1,…, *I_j,k_*) are independent and identically distributed (i.i.d.), following a normal distribution given by

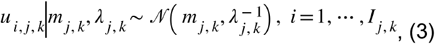

where the precision *λ_j,k_* follows a Gamma distribution, i.e.,

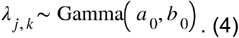

Moreover, to characterize the shift of the expectation in Eq. (3) from the background due to its functional effect, we model *mj,k* by a three-component Gaussian mixture model, i.e.,

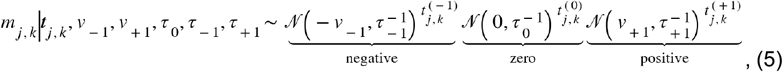

where the precisions follow

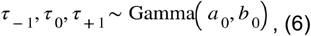

and *v*_-1_ and *v*_+1_ are non-negative variables quantifying the absolute values of effect size shifts for the negative and positive components, respectively.

To impose non-negativity over *v*_-1_ and *v*_+1_, here we employ the rectification nonlinearity technique proposed previously (Harva and Kabán, 2007). In particular, we assume *v*_-1_ and *v*_+1_ follow

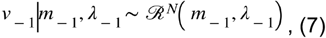

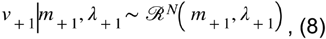

in which the rectified Gaussian distribution is defined via a dumb variable. Specifically, we first define *v*_-1_ and *v*_+1_ by

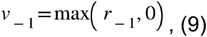

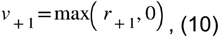

which guarantee that *v*_-1_ and *v*_+1_ are non-negative. The dump variable *r*_-1_ and *r*_+1_ follow Gaussian distributions given by

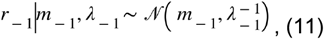

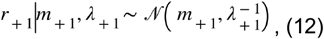

where *m*_±_ and *λ*_±_ follow the Gaussian-Gamma distributions, i.e.,

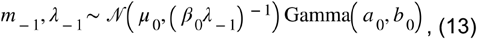

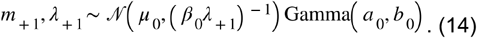

The indicator variables in Eq. (5) denote whether that region is ALS-associated or not. Indeed, we define the region to be disease-associated if 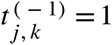 or 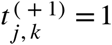, and to be non-associated otherwise. To simplify the analysis, we put a symmetry over 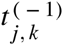 and 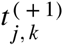, and define the distribution by

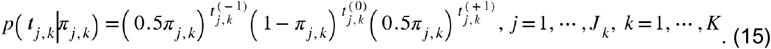

Furthermore, the probability parameter *π_j,k_* in Eq. (15) is given by

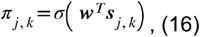

where *σ*(·) is the sigmoid function, ***s**_j,k_* is the vector of epigenetic features for the *j*-th region in the *k*-th LD block, and the weight vector ***w*** follows a multivariate normal distribution, i.e.,

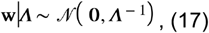

and *Λ* follows

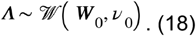

In our study, the epigenetic features ***s**_j,k_* were calculated as the overlapping ratios of that region with the narrow peaks of ATAC-seq and histone ChIP-seq, respectively.

Based on Eqs. (1) to (18), we are interested in calculating *p*(***T | Z, S***) wherein the calculation of integrals is intractable. Here we seek for approximate inference based on the mean-field variational inference (MFVI)(Blei et al., 2017). To reduce false positives, we set a hard threshold for 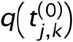 with respect to the ATAC-seq signal, where we set 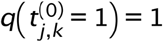 if the corresponding region overlaps no ATAC-seq peak. This was motivated by our particular interest in active regions. More technical details, including a coordinate ascent-based inference algorithm, were provided in Supplementary Notes.

In this study, we ran the inference algorithm per chrompsprpe to accelerate the computation. The *Q*^+^- and *Q*^−^-scores were defined as *q*(*t*^(+1)^ =1) and *q*(*t*^(+1)^ =1), respectively, and we also defined the *Q*-score as *Q*=*Q*^+^+*Q*^−^. To prioritize RefMap-scored regions, we set a cutoff of 0.95 and defined those regions with either *Q^+^*- or *Q^−^*-score larger than the cutoff as significant regions (i.e., ALS-associated regions) (**Supplementary Table 5**).

Code relevant to RefMap is available on request.

#### Target gene identification

After identifying ALS-associated regions based on RefMap, we mapped those active regions to their target genes for a better understanding of their functions. In particular, we performed such mapping according to to two principles: (i) assign to a gene if the region overlaps the gene or the region up to 10kb either side of the gene body; (ii) assign to a gene if the region overlaps a loop anchor harboring the transcription start site (TSS) of that gene. The loops were called from the Hi-C data sequenced from the iPSC-derived MNs. Note the only transcripts with TPM>=1 were kept for downstream analysis.

#### Network analysis

We first downloaded the human PPIs from STRING v11, including 19,567 proteins and 11,759,455 protein interactions. To eliminate the bias caused by hub proteins, we first carried out the random walk with restart algorithm(Wang et al., 2015) over the PPI network, wherein the restart probability was set to 0.5, resulting in a smoothed network after preserving the top 5% predicted edges. To decompose the network into different subnetworks/modules, we performed the widely-used Louvain algorithm(Blondel et al., 2008b), a classic community detection algorithm that searches for densely connected modules by optimizing the modularity. After the algorithm converged, we obtained 912 modules with an average size of 18.39 nodes (**Supplementary Table 11**). Two modules (M421 and M604) were significantly enriched (FDR<0.1) with our RefMap genes based on the hypergeometric test followed by the BH correction (**Supplementary Table 11**).

To test whether the network modularity could be observed by chance, we randomly shuffled the edges of the network while preserving the number of neighbors for each node (Milo et al., 2002). We generated 100 such randomized networks followed by the Louvain decomposition, against which the modularity of the smoothed PPI network was tested.

#### Rare variant burden tests

ALS features a polygenic rare variant architecture (van Rheenen et al., 2016), therefore, all searches for pathogenic variants in enhancer and coding regions featured a filter for MAF within the Genome Aggregation Database (gnomAD) of <1/100 control alleles (Lek et al., 2016). Additional filtering varied reflecting differences in function between enhancer, promoter and coding sequence. In enhancer regions, variants were included only if evolutionary conserved based on a LINSIGHT score >0.8 (Huang et al., 2017). We also utilized an independently compiled score for ALS-associated regulatory variation (Chen et al., 2016a): variants were excluded with a DIVAN score <0.5. In promoter regions, we utilized two independent scores for functionality and pathogenicity: variants were included in burden testing if their CADD (Rentzsch et al., 2019) score >25 and GWAVA (Ritchie et al., 2014) score >0.5. In coding regions, we filtered for variants with impact on protein function as defined by snpeff (Cingolani et al., 2012): variants annotated HIGH/MODERATE/LOW impact were included, but we excluded variants annotated ‘synonymous’ or ‘TF_binding_site_variant’ because these functions are independent of amino acid sequence.

The optimal unified test (SKAT-O) was used to perform burden testing in enhancer and promoter regions because it is optimized for large numbers of samples and for regions where a significant number of variants may not be causal (Lee et al., 2012). SKAT tests upweight significance of rare variants according to a beta density function of MAF in which *W_j_* = *Beta*(*p_j_*, *a*_1_, *a*_2_), where *p_j_* is the estimated MAF for SNP_*j*_ using all cases and controls, parameters *a*_1_ and *a*_2_ are prespecified, and *a*_2_=2500 was chosen for all statistical tests. To adjust for confounders including population structure, burden testing used the first ten eigenvectors generated by principal components analysis of common variant profiles, sequencing platform and sex as covariates.

#### Morphological assessment of differentiated SH-SY5Y cells

To confirm neuronal differentiation and to assess for changes consistent with axonopathy, semi-automated analysis of neurite length was performed using the SimpleNeuriteTracer plugin for FIJI (Longair et al., 2011). 2D images were converted to 8-bit grayscale and successive points along the midline of a neural process were selected. The software automatically identified the path between the two points. Tracing accuracy was improved using Hessian-based analysis of image curvatures. The AnalyzeSkeleton plugin (Arganda-Carreras et al., 2010) was used to quantify the morphology of the traces including the length of neurites. In the case of joined neurites the shorter path length was assigned to ‘branches’. To determine whether observed changes in neurite length are significant three fields of view were analyzed and differences were assessed by a Student’s t-test, where a one-tailed test was chosen based on the hypothesis that ALS-associated mutations would reduce neurite length.

#### Quantitative PCR and MTT assays

Relative mRNA expression values were then calculated using the 2^−ΔΔCT^ method (Schmittgen and Livak, 2008). Statistical analysis was conducted in GraphPad Prism 7 (La Jolla, CA) and R (v4.0.2). All bar graphs show the mean ± SD. To identify statistical differences between treatment groups utilised Student’s unpaired t-test.

## Supplemental Notes for

### 1 Mathematical foundation of RefMap

Here, we provide a mathematical theory to justify Eq. 1 in the Method section of the main text. To facilitate the development of the theory, we first describe a universal discriminative framework that models the relationship between the genotype and phenotype, and then deduce a general distribution over summary statistics from this framework. Based on this result, a flexible probabilistic model that characterizes summary statistics with various prior structures can be developed, which generalizes multiple previous studies [1–4, 6–8]. In particular, Equation 1 of `RefMap` follows directly after assuming a linear relation between the genotype and phenotype. In the following, we will develop the framework in both cases of quantitative trait and case-control studies.

#### 1.1 Quantitative trait studies

We start from considering a general genotype-phenotype model for continuous traits, i.e.,

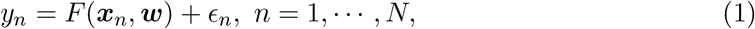

in which *N* is the sample size, ***x**_n_* and *y_n_* are the genotypes and phenotype for the *n*th sample, respectively, *F* is an unknown (usually non-linear) function with parameters ***w*** determining personal phenotype from his/her genotypes, and *ϵ_n_* is the random noise following

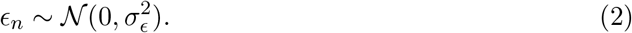

Note that as a routine procedure, genotypes are first standardized by

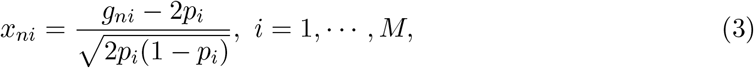

where *M* is the number of alleles, *g_ni_* is the genotype of the *i*th allele for the *n*th sample, and *p_i_* is the frequency of the ith allele in the study cohort. After standardization, the sample mean and sample variance of each allele are 0 and 1, respectively. Moreover, we adopt a general setting and treat both genotypes and function parameters as random variables, yielding

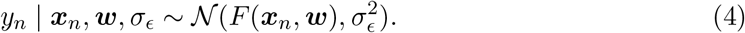

Following the conventional annotation in the genome-wide association study (GWAS), the estimated effect sizes 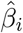 for individual alleles are the most widely-used summary statistics, which are closely related to *χ*^2^ and *Z*-score. Given the genotype standardization, we have

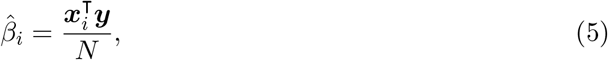

where ***x**_i_* is the genotype vector for the ith allele and ***y*** = *y*_1:*N*_. With matrix representation, we have

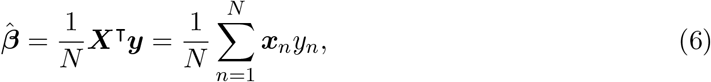

where ***X*** = (*x_ni_*)∈ ℝ^*N×M*^. Indeed, we have the following theorem characterizing the asymptotic distribution of 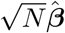.

##### Theorem 1.

*Given the definitions in Eqs. 1, 2 and 5, when the sample size N is large enough, we have*

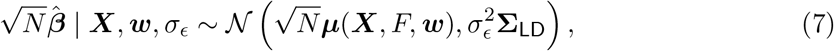

*where* **Σ_LD_** *is the in-sample linkage disequilibrium (LD) matrix quantifying SNP correlations, and **μ**(**X, F, w**) is a quantity depending on the genotypes and the discriminative function F*.

*Proof.* We first show tha 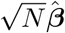 follows a normal distribution asymptotically. In fact, according to Eq. 6, given the genotypes and the discriminative function, 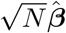 can be computed by the sum of ***x**_n_y_n_*, which are independent with each other but with different expectations. On the other hand, the variance of ***x**_n_y_n_* is given by

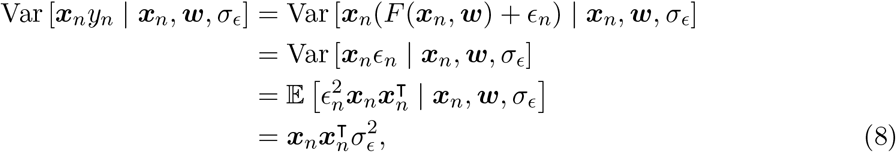

yielding

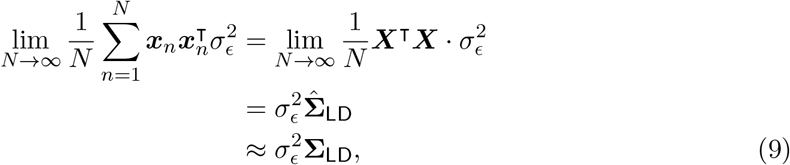

in which the estimated LD matrix 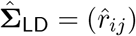 is given by

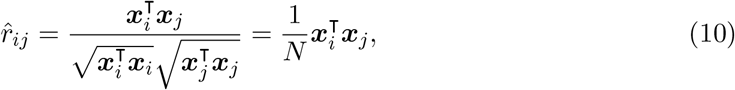

and the last approximation is guaranteed by 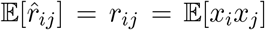. Therefore, according to the multivariate Lindeberg-Feller central limit theorem (CLT), we conclude that 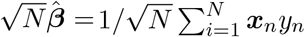 asymptotically follows a normal distribution with covariance 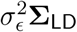, whose expectation is given by

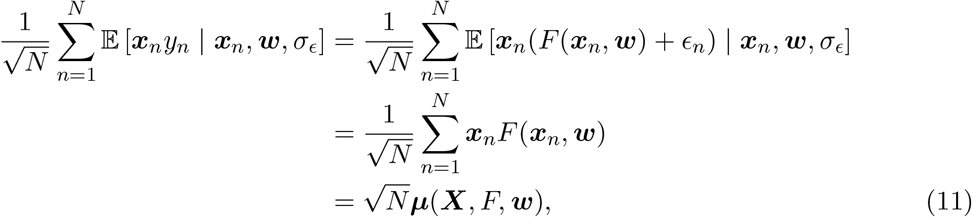

where ***μ***(·) is defined as

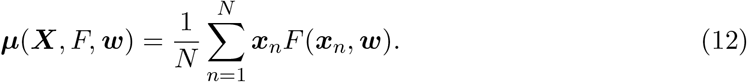

This completes the proof.

Note that if we use *Z*-scores computed by GWAS as the approximation of 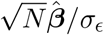, i.e., dividing 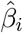 by its estimated standard error, we have

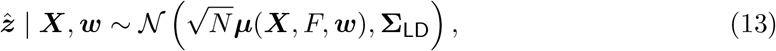

in which σy is absorbed into μ(o) for annotation brevity.

#### 1.2 Case-control studies

We state the analysis for case-control studies using a Bernoulli distribution over case-control status, i.e.,

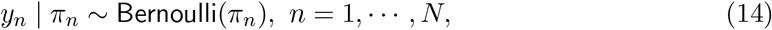

whose logit is defined similarly as Eq. 1 but without random noise, i.e.,

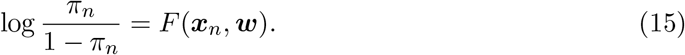

After a few calculations we can easily get

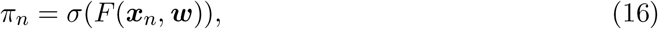

where *σ*(·) is the sigmoid function defined by *σ*(*x*) = 1/(1 + exp(-*x*)).

To facilitate the following analysis, here we illustrate the standardization procedure in more detail, i.e.,

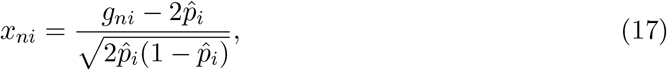

where *g_ni_* is the genotype coded by 0, 1 and 2, and 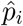 is the in-sample allele frequency. Therefore, suppose we have the same number (*N*/2) of cases and controls in the study cohort, the widely-used *Z*-scores for case-control studies defined as

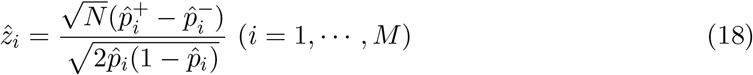

can be written as

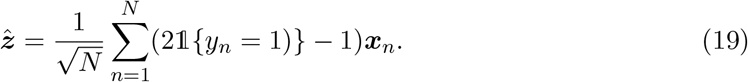

Again, utilizing the multivariate Lindeberg-Feller CLT, we can derive the asymptotic conditional distribution of 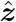, which is approximately the same as that in the quantitative trait studies (Eq. 13). In particular, we have the following result.

##### Theorem 2.

*Given the definitions in Eqs. 14, 15 and 19, when the sample size N is large enough, we have*

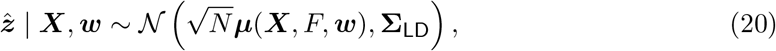

*where* **Σ_LD_** *is the in-sample LD matrix, and **μ**(**X, F, w**) is a quantity depending on the genotypes and the discriminative function*.

*Proof.* Conditioned on **X** and **w**, the variance of 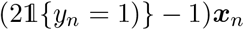 can be calculated as

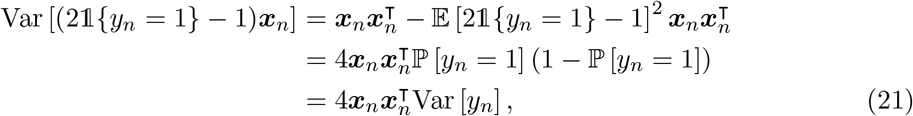

where the conditions are omitted for brevity. In fact, as Var [*y_n_* | ***x**_n_, **w***] < 1, we conclude that the average of variance 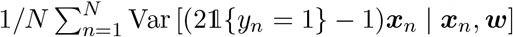 converges as *N* ! ∞, whose limit is denoted as **Σ**_∞_. According to the multivariate Lindeberg-Feller CLT, the asymptotic conditional distribution of 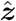 is a normal distribution with covariance matrix **Σ**_∞_.

To get a clearer structure of **Σ**_∞_, we now apply a few approximations for Eq. 21. In particular, we have

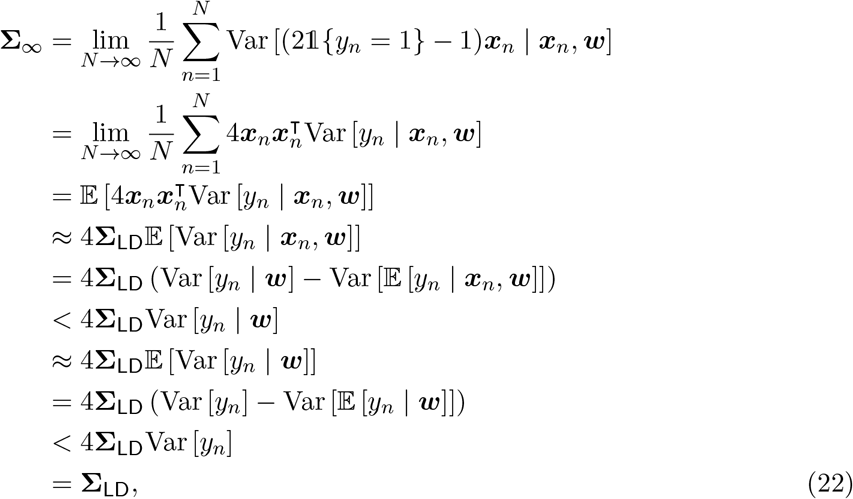

in which the third and the fourth “=” come from the law of total variance, the first and the second “<” are implied by the positivity of variance, and **Σ_LD_** is the in-sample LD matrix. For the last “=”, we argue that the expectation and variance in Eq. 22 are taken over the sampling space in case-control studies, rather than the general population. Under the assumption of equal number of cases and controls, the sampling disease prevalence is 0.5, yielding Var [*y_n_*] = 0.25.

Furthermore, the expectation of the asymptotic conditional distribution can be calculated as

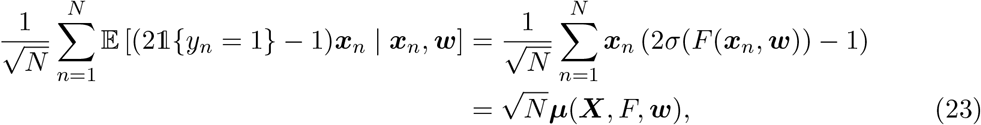

where we define

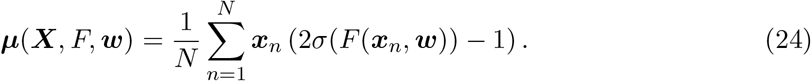

This completes the proof.

#### 1.3 A linear model for RefMap

We consider a linear model that underlies the design of `RefMap`. Specifically, in the quantitative trait studies, we define

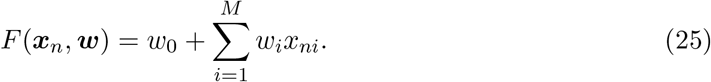

Note that this linear model has been widely used in traditional GWAS studies [1—3], and *w_i_* is called the *effect size* of the ith allele. The linear model for case-control studies can be developed similarly by considering the approximation of sigmoid function using its Taylor expansion. Therefore, the expectation of the asymptotic distribution of *Z*-scores can be calculated as

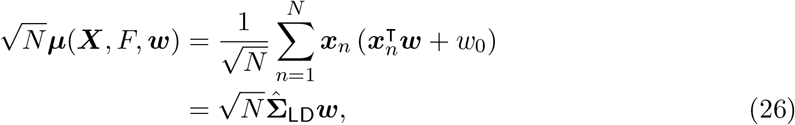

indicating that the expected *Z*-score for each allele is determined by its effect size as well as its strongly-associated neighbors. By absorbing 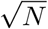 into *w*, we eventually get Eq. 1 in the `RefMap` model.

### 2 Inference for RefMap

The RefMap model was defined in Eqs. 1 to 18 in the Method section of the main text. Here, we are interested in the posterior *p*(***T*** | ***Z, S***), whose exact calculation is intractable. Therefore, we seek for approximate inference based on the mean-field variational inference (MFVI). Basically, we first assume that the approximate posterior over latent variables factorizes, indicating conditional independence across latent variables, and then perform approximate inference by optimizing the *evidence lower bound* (ELBO) with respect to factorized proposal distributions, i.e.,

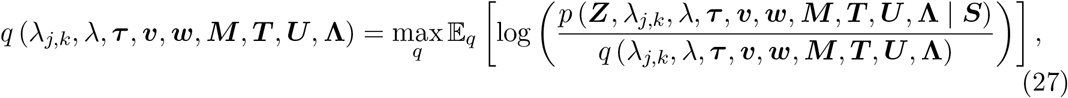

which can be shown to be equivalent to minimizing the Kullback-Leibler (KL) divergence between the true posterior and its proposal.

In the following, we will first introduce several specific techniques we used in MFVI, and then summarize the update rules for different variational parameters. At last, a coordinate ascent-based VI algorithm will be given.

#### 2.1 Rectification nonlinearity

We impose non-negativity on *v*_−1_ and *v*_+1_ using the technique of rectification nonlinearity proposed in [5]. This technique relaxes the sparsity constraint over factors and meanwhile enjoys tractable variational inference.

We first note that the approximate posterior *q*(*r*_−1_) from MFVI follows the free-form solution

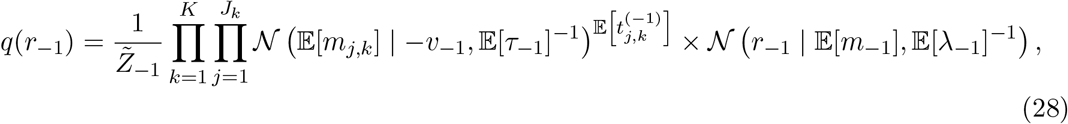

where 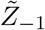 is the normalization term to be computed later. Moreover, it can be easily shown that Eq. 28 can be written as *q*(*r*_−1_) = *q_p_*(*r*_−1_) + *q_n_*(*r*_−1_) with the form

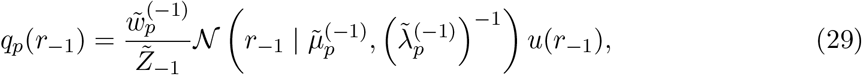

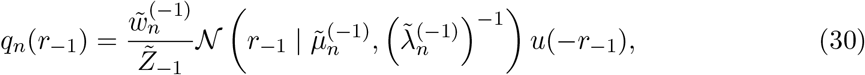

in which

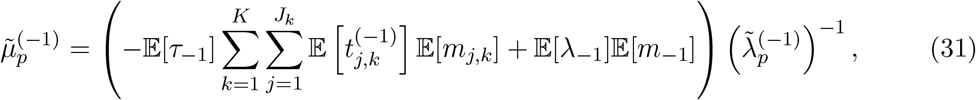

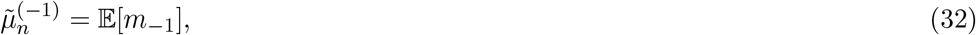

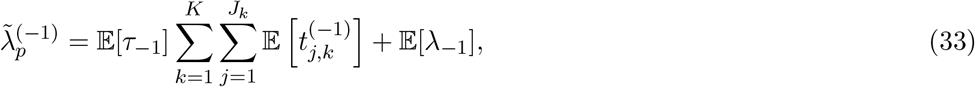

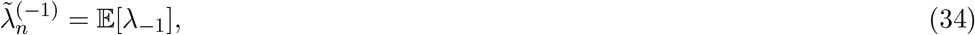

and *u*(·) is the standard step function. With Eqs. 31 to 34, 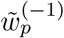 and 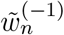 can be computed by integrating Eqs. 28, 29 and 30 with respect to *r*_−1_. Then the normalization term is given by

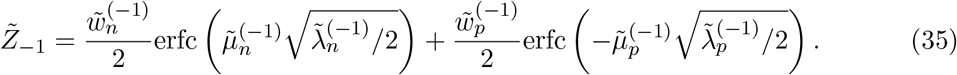

The moments for posteriors are obtained by

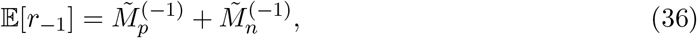

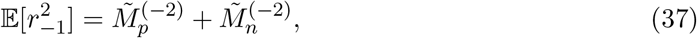

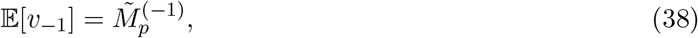

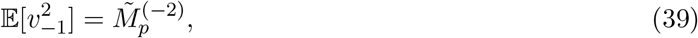

where

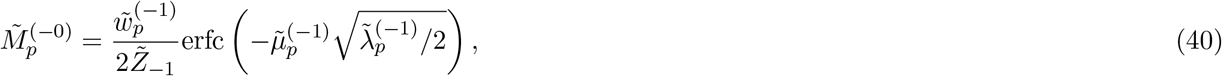

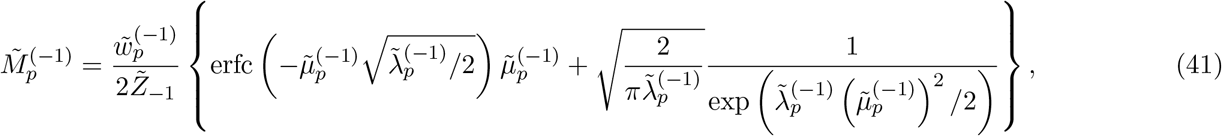

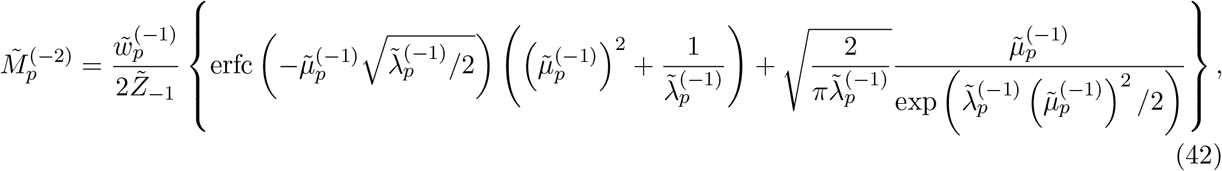

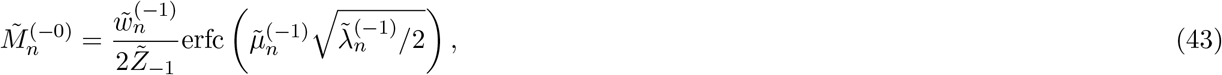

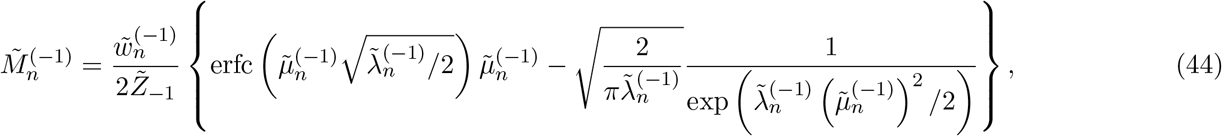

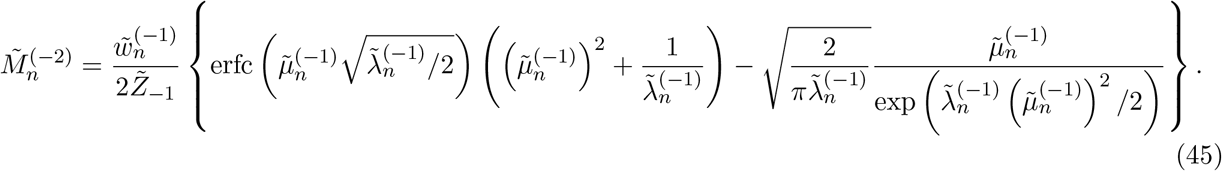

Similar to *q*(*r*_−1_), the posterior *q*(*r*_+1_) also follows a free-form solution given by

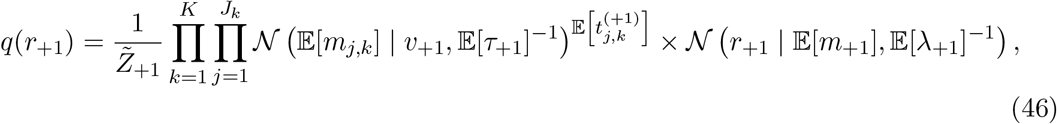

where 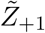 is the normalization term. Equation 46 can also be written as *q*(*r*_+1_) = *q_p_*(*r*_+1_) + *q_n_*(*r*_+1_) with the form

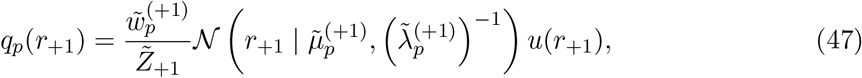

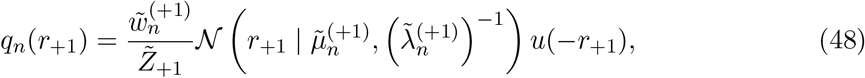

in which

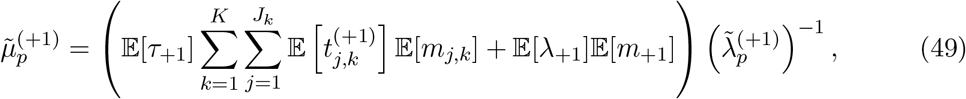

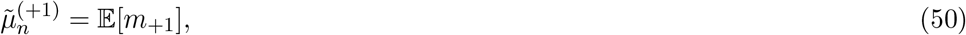

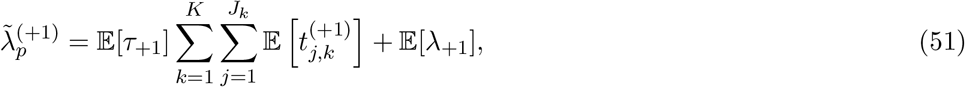

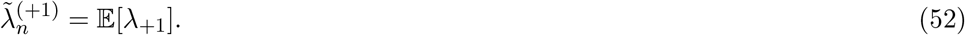

After computing 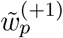 and 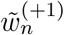, the normalization term is given by

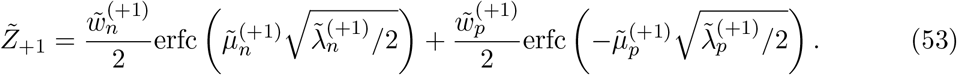

The moments for posteriors are obtained by

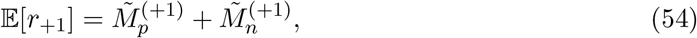

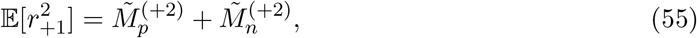

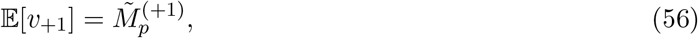

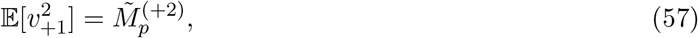

in which

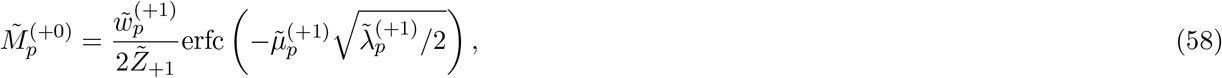

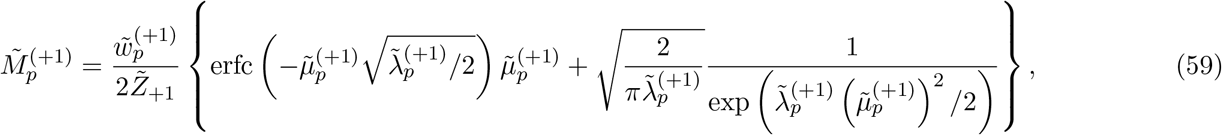

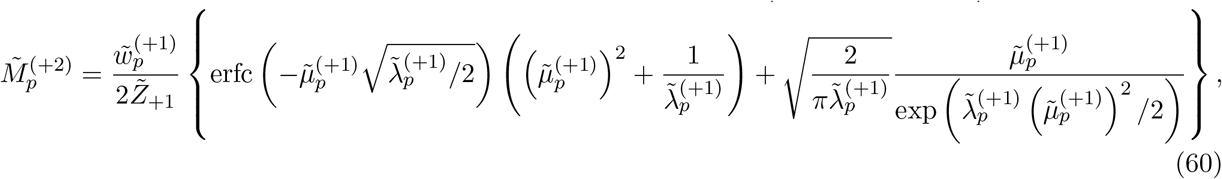

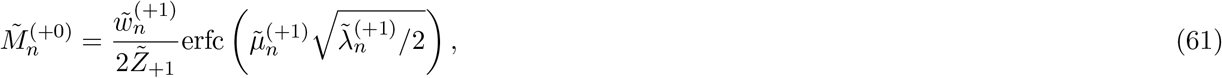

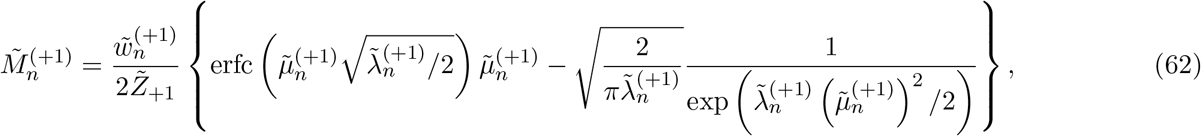

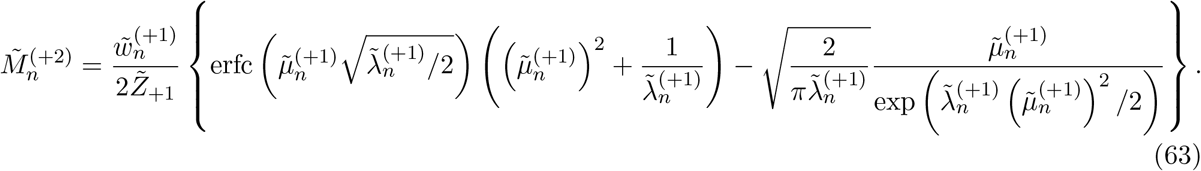

#### 2.2 Local variational method

We adopt the local variational method to tackle the intractability of MFVI for ***w*** due to the introduction of the sigmoid function (Eq. 16 in the Method section). In particular, we have the following result regarding Eq. 15 in the Method section:

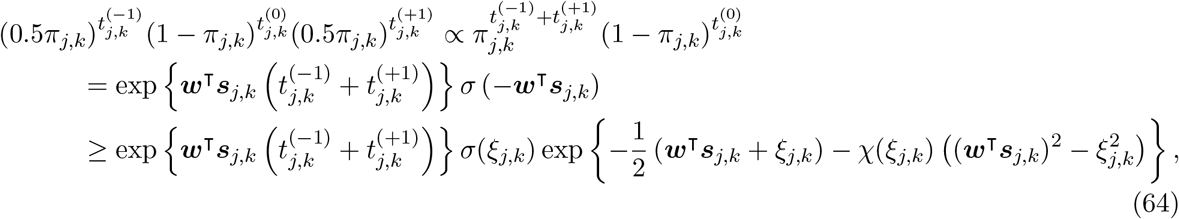

where

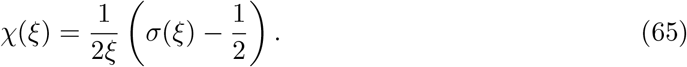

Then we can perform standard MFVI with respect to the lower bound of Eq. 64, which yields

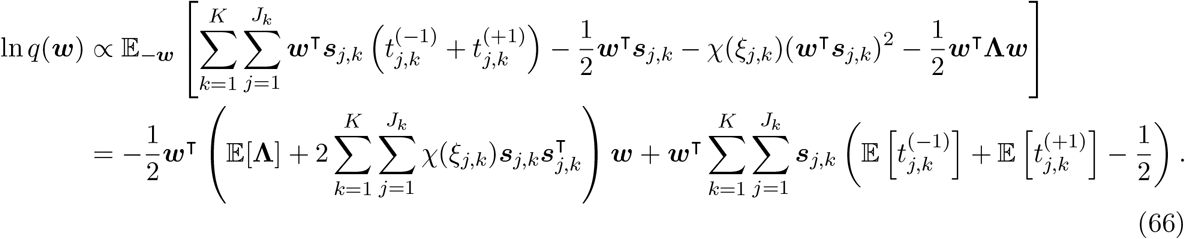

This indicates that *q*(***w***) follows a normal distribution given by

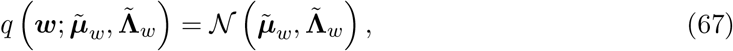

in which

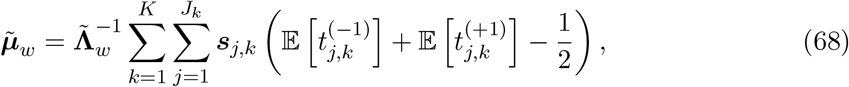

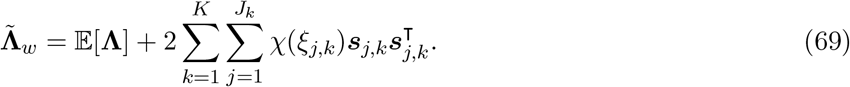

#### 2.3 Update rules for other variational parameters

For other latent variables in `RefMap` besides *v*_−1_, *v*_+1_ and ***w***, we carry out the naive MFVI and obtain

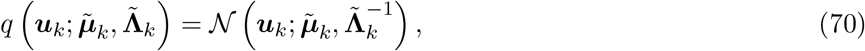

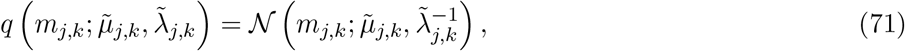

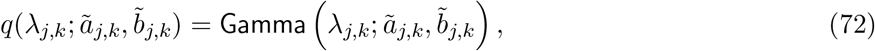

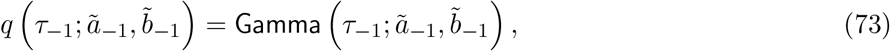

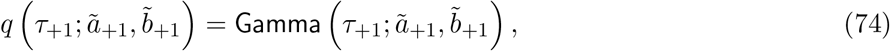

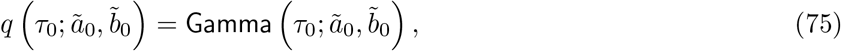

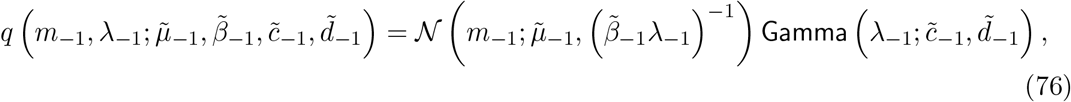

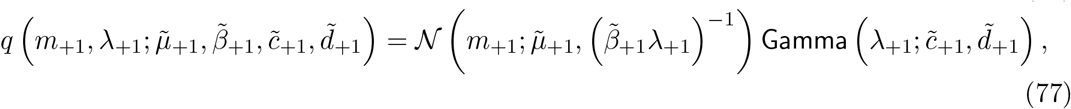

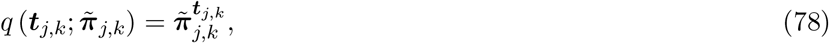

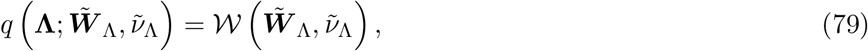

in which

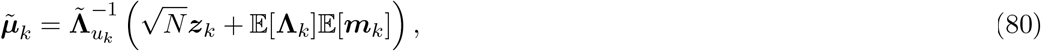

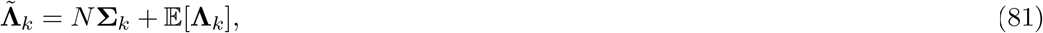

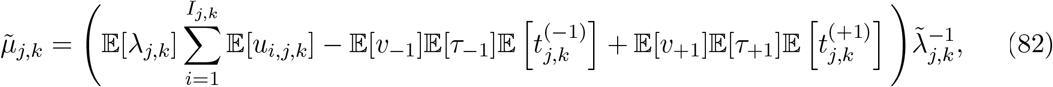

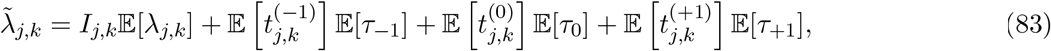

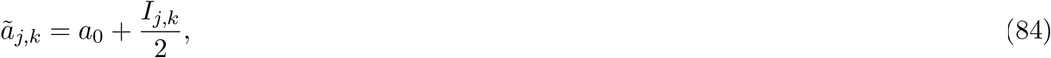

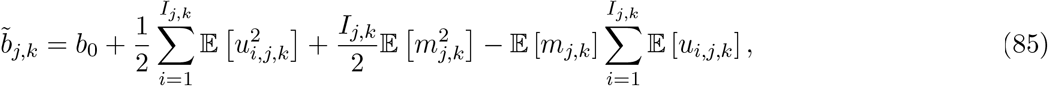

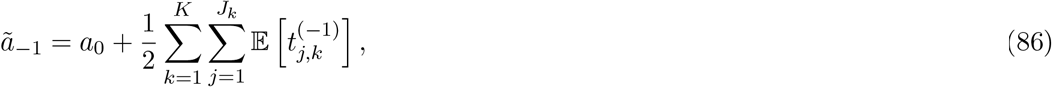

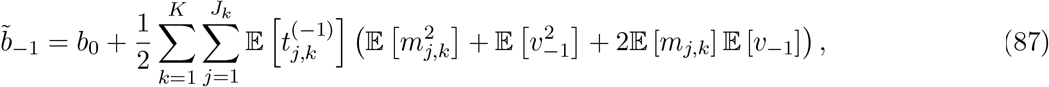

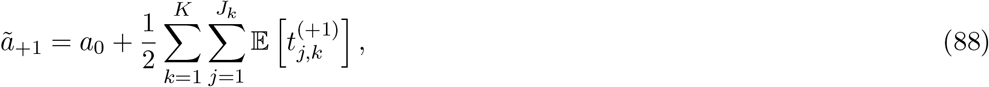

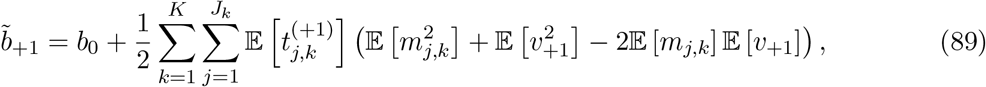

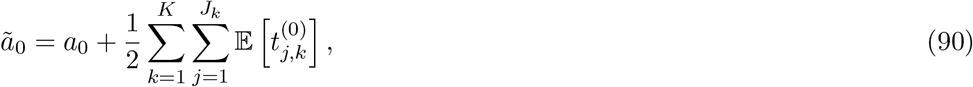

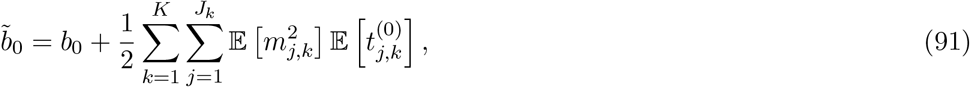

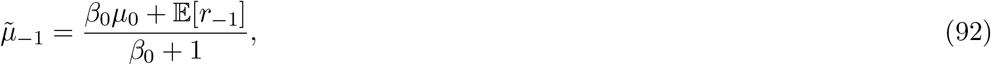

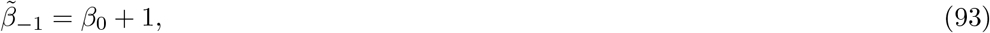

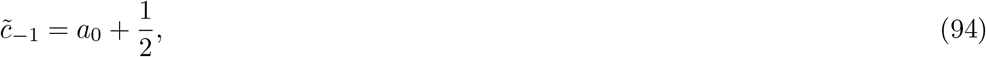

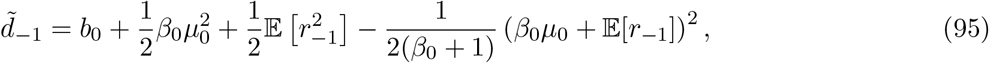

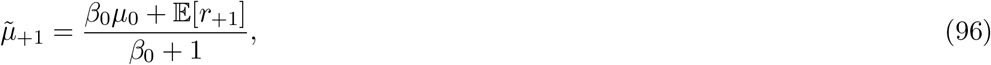

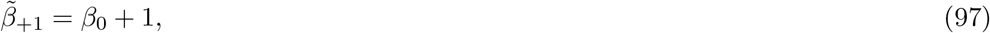

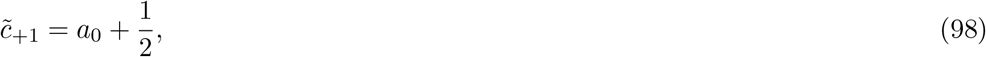

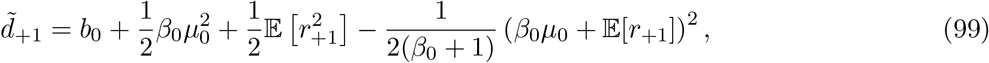

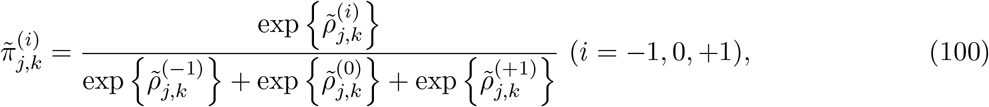

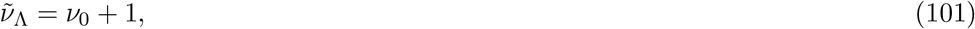

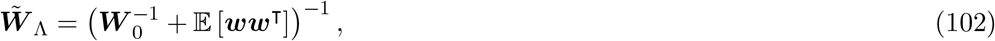

and we define

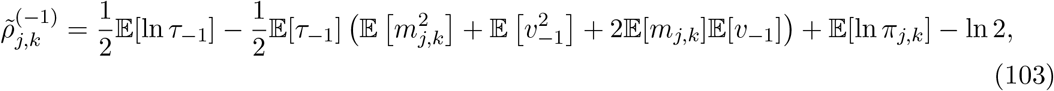

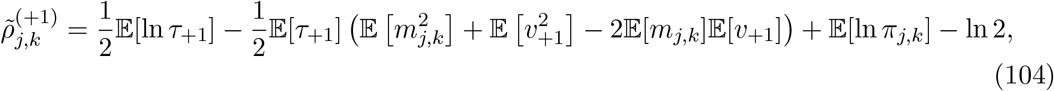

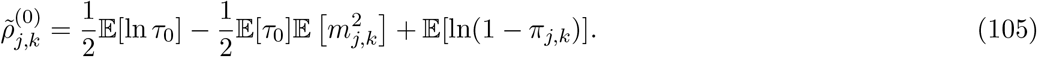

**Algorithm 1:**
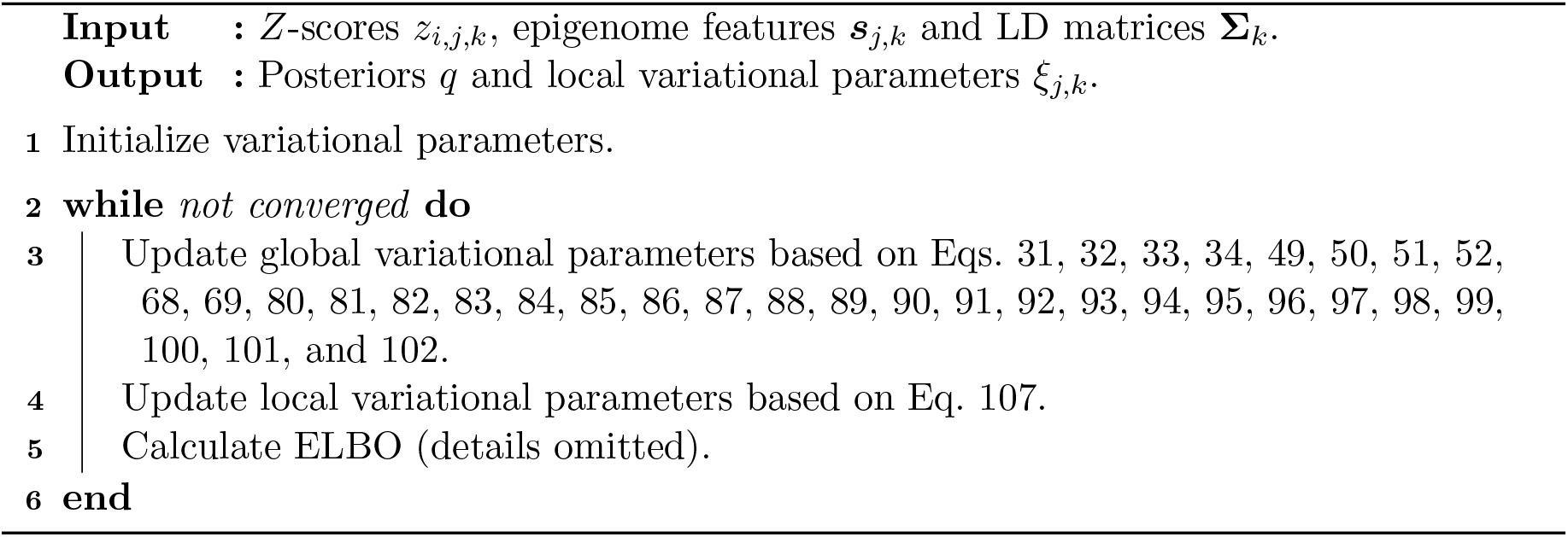
MFVI for RefMap

#### 2.4 Update rules for local variational parameters

One needs to maximize the lower bound on marginal likelihood in Eq. 64 with respect to *ξ_j,k_* to rationalize the local variational inference. In particular, we have the following optimization problem

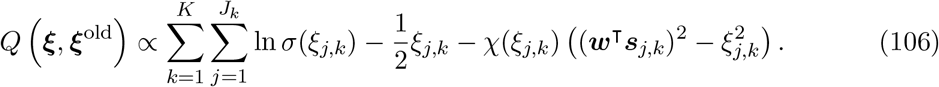

Solving the above problem with respect to each *ξ_j,k_* gives its update rule

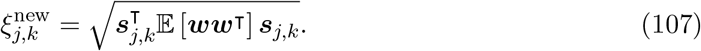

#### 2.5 Coordinate ascent algorithm for MFVI

With the above update rules we can construct a coordinate ascent algorithm to update variational parameters iteratively until convergence (i.e., the change of ELBO falls below a threshold which was set to be 10^−6^ in our study). The inference algorithm is summarized in Algorithm 1.

